# Rapamycin Immunomodulation Utilizes Time-Dependent Alterations of Lymph Node Architecture, Leukocyte Trafficking, and Gut Microbiome

**DOI:** 10.1101/2024.10.01.616121

**Authors:** Long Wu, Allison Kensiski, Samuel J Gavzy, Hnin Wai Lwin, Yang Song, Michael France, Ram Lakhan, Dejun Kong, Lushen Li, Vikas Saxena, Wenji Piao, Marina W. Shirkey, Valeria Mas, Bing Ma, Jonathan S Bromberg

**Affiliations:** University of Maryland School of Medicine, Department of Surgery, Baltimore, MD, USA; University of Maryland School of Medicine, Center for Vascular and Inflammatory Diseases; University of Maryland School of Medicine, Institute for Genome Sciences, Baltimore, MD, USA; Department of Microbiology and Immunology, University of Maryland School of Medicine, Baltimore, MD, USA

**Author notes:** Corresponding Authors: Jonathan S. Bromberg, MD, PhD, Bing Ma, PhD.

## Abstract

Transplant recipients require lifelong, multimodal immunosuppression to prevent rejection by reducing alloreactive immunity. Rapamycin, a mechanistic target of rapamycin (mTOR) inhibitor, is known to modulate adaptive and innate immunity, while the full spectrum of its immunosuppressive mechanisms remains incompletely understood. Given the broad expression of mTOR, we investigated the understudied effects of rapamycin on lymph node (LN) architecture, leukocyte trafficking, and the gut microbiome and metabolism after 3, 7, and 30 days of rapamycin treatment, to characterize the early, intermediate, and late changes. Rapamycin significantly reduced CD4+ T cells, CD8+ T cells, and regulatory T (Treg) cells in peripheral LNs, mesenteric LNs, and the spleen over time. Rapamycin induced early pro-inflammation transition to pro-tolerogenic status, by modulating the LN laminin α4:α5 expression ratios through LN stromal cells laminin α5 expression and by adjusting Treg numbers and distribution. Additionally, rapamycin significantly altered gut microbiota composition and metabolic functions, shifting the Bacteroides to Firmicutes ratio and increasing amino acid bioavailability in the gut lumen. These effects were evident by 7 days and became most pronounced by 30 days in naïve mice, with notable changes as early as 3 days in allogeneic splenocyte-stimulated mice. These findings reveal a novel mechanism of rapamycin’s action through time-dependent modulation of LN architecture and gut microbiome, which orchestrates changes in immune cell trafficking, providing a new framework for understanding and optimizing immunosuppressive therapies.

## INTRODUCTION

Prevention of solid organ transplant rejection requires the use of lifelong, multimodal immunosuppression to dampen both adaptive and innate alloreactive immunity. One major mainstay is rapamycin (aka. sirolimus), a bacterial fermentation product recognized for its immunosuppressive and antiproliferative properties through the inhibition of the mechanistic target of rapamycin (mTOR), a critical kinase in cell cycle regulation and immune response modulation (1, 2). The mTOR pathway is a key regulator of cellular metabolism and growth, influencing nutrient availability, growth factors, and stress responses. Rapamycin has been shown to promote autophagy, delay cellular senescence, and act as an anti-proliferative agent in certain cancers (3, 4). The anti-proliferative effects are beneficial in calcineurin inhibitor-free regimens, preserving renal function and reducing the incidence of post-transplant cancers (5), although these same properties can lead to side effects such as increased infection risk, impaired wound healing, and metabolic disturbances (6). Studies have also highlighted rapamycin’s protective effects against atherosclerosis and its successful use in patients with coronary artery disease (7). Despite its established role in mTOR inhibition, the impact of rapamycin on other immune populations and its precise mode of action has not been fully investigated. Given its broad impact on immune landscape, it is critical to understand its molecular mechanisms for optimizing its clinical use and enhancing longevity in solid organ transplant patients.

Rapamycin targets mTOR complex 1 (mTORC1) and mTOR complex 2 (mTORC2) and blocks the activation and proliferation of T cells and B cells in response to cytokine stimulation to exert immunosuppressive effects (3, 4, 8). Rapamycin modulates various signaling pathways, promoting the maintenance of memory T cells and improving their ability to respond to subsequent exposures to the same antigen (9, 10). It also preserves and supports the maturation-resistance and tolerogenic properties of dendritic cells (DC) (11, 12). Beyond its immunosuppressive properties, rapamycin’s unique attributes extend to cancer therapy. The mTOR axis also regulates the expression of programmed death-ligand 1 (PD-L1) in cancer cells, downregulating PD-L1 expression and thereby enhancing the effectiveness of immune checkpoint inhibitors in cancer therapy (13, 14). Rapamycin modulates innate immunity by promoting an anti-inflammatory phenotype in DCs via enhanced mitochondrial metabolism and increased IL-10 production (8). Recent studies have demonstrated its ability to delay the onset of cellular senescence and regulate metabolic pathways similar to caloric restriction, interventions known to extend lifespan in various species (3). Rapamycin’s regulation of autophagy, a process required for maintaining amino acid levels and protein synthesis during nitrogen starvation, can significantly influence amino acid balance and related cellular metabolism and immune responses (15).

Beyond their primary action on immune cell populations, immunosuppressants can also significantly affect the gut microbiome, the largest immune organ in the human body, thus influencing alloimmunity and graft survival (16–20). In this way, immunosuppressants elicit secondary effects on both local and systemic immune responses, which in turn can influence allograft outcomes (21–25). Recent work by us and others revealed a complex interplay between the commensal gut microbiome, immunosuppressant therapies, and the host immune system (22, 26–29). These investigations highlight that immunosuppressant-induced changes in the gut microbiome, alongside the use of antimicrobials, can impact overall immune homeostasis. Immunosuppresants primarily affect anaerobic bacteria, including *Ruminococcaceae*, *Lachnospiraceae*, Firmicutes, *Bacteroides*, and Clostridiales (30–32). Furthermore, immunosuppressants use has been associated with an increase in colonization of uropathogenic *Escherichia coli* and *Enterococcus sp.* (25, 33, 34). Overall the impact of immunosuppressants extends beyond their direct effects on immune cells to include significant modulation of the gut microbiome. However, the precise alterations in microbiome composition and function induced by rapamycin, along with their underlying mechanisms, remain to be elucidated.

In addition to multi-directional gut microbiome-endothelium-immune cell interactions, rapamycin’s impact on other lymphoid organs, particularly lymph nodes (LNs), remains poorly understood. LN structure relies on LN stromal cells (LNSCs) to regulate the position and interaction of lymphocytes with antigen-presenting cells via chemokines, cytokines, and stromal fibers (35–37). LNSCs, which include fibroblastic reticular cells (FRCs), lymphatic endothelial cells (LECs), and blood endothelial cells (BECs), form a critical infrastructure that supports immune cell trafficking and interactions within the LN microenvironment. FRC-derived laminins play a crucial role in modulating global immune states by balancing tolerance and immunity (38). Laminins, as extracellular matrix proteins, influence the structural integrity of the LN and impact immune cell behavior (38). Changes in LN architecture driven by laminins can be induced by various factors, including the gut microbiota and alloimmunity (28). For example, the gut microbiota can influence LN structure through microbial metabolites and immune signaling (28). Similarly, alloimmunity can lead to structural changes in the LN, affecting immune tolerance and graft survival (39). Additionally, ischemia reperfusion injury of the graft during transplantation can disrupt LN structure and FRC function, leading to immunologic scarring (40, 41). Pathologic alterations to this cellular network pose significant challenges for maintaining immune homeostasis and preventing chronic graft rejection. Investigating rapamycin’s effect on LN architecture and LNSCs offers key insights into its immunosuppression mechanisms.

This study investigates the contribution of rapamycin to LN architecture, leukocyte trafficking, the gut microbiome, and the mechanisms underlying these changes. We examined microanatomic cell positioning and interactions, as well as LN stromal fiber structure, building on our previous work that demonstrated the importance of architectural and cellular changes within the LN cortex, including the cortical ridge (CR) and around the high endothelial venules (HEV), in mediating immune tolerance and suppression (38, 42). To capture the early, intermediate, and late effects of rapamycin, we performed temporal characterization after 3, 7, and 30 days of treatment. We revealed dynamic, time-dependent effects of rapamycin on LN architecture and Treg localization, showing a transition from early pro-inflammatory changes to a later pro-tolerogenic environment. This transition was mediated by the modulation of the laminin α4:α5 ratio in LN stromal cells via altering laminin α5 expression, with a higher laminin α4:α5 ratio indicative of a tolerogenic environment. These observations were validated across transgenic mouse lines and models, including naïve, laminin α4 knockout, laminin α5 knockout, and allogeneic splenocyte-stimulated mice. Further, rapamycin significantly altered gut microbiota composition, intestinal Tregs and metabolic functions. These effects were evident by 7 days and became most pronounced by 30 days in naïve mice, emphasizing the incremental and dynamic effect of rapamycin. Our study reveals rapamycin’s multifaceted, time-dependent effects on immune regulation via LN architecture and gut microbiome, offering a new lens for optimizing long-term immunosuppressive therapies to enhance graft survival and patient longevity.

## RESULTS

### Rapamycin reduces lymphocytes in LNs and spleen

To investigate the immunological effects of rapamycin within secondary lymphoid organs, C57BL/6 mice received rapamycin (5 mg/kg/day i.p.) (43). The mice were characterized after 3-, 7-, and 30-days of treatment representing early, intermediate, and late changes. We assessed cell populations in peripheral lymph nodes (pLNs), mesenteric lymph nodes (mLNs), and spleen, focusing on CD4+ and CD8+ T lymphocytes and Tregs via flow cytometry. Significant reductions in pLNs weight and overall cell counts were observed, including inguinal, axillary, brachial, and cervical LNs at all three time points, consistent with rapamycin’s known inhibitory effect on cell proliferation and metabolism **(Supplemental Figure 1A-D)**. Rapamycin reduced specific immune cell populations within the pLNs, including CD4+ and CD8+ T lymphocytes and Foxp3+ Tregs at all time points **(Figure 1A, D)**, demonstrating significant and sustained immunomodulatory effects. In the mLNs, rapamycin significantly reduced CD4+ and CD8+ T lymphocytes and Foxp3+ Tregs after 7 and 30 days, but not on day 3 **(Figure 1B, D)**. In the spleen, similar decreases in CD4+ T cells and Foxp3+ Tregs were noted on days 7 and 30, with a decline in CD8+ T cells observed only on day 7 **(Figure 1C, D)**. Rapamycin decreased the Treg/non-Treg ratio in the spleen but not in pLNs and mLNs (**Supplemental Figure 1E-G**), suggesting an uneven impact on various T cell populations. Overall, these data demonstrate that rapamycin significantly reduces lymphocytes and Treg cell counts in LNs and spleen, a manifestation of its immune suppressive effect. Notably, rapamycin showed more pronounced effects on CD4+ and CD8+ T lymphocytes and Treg cells in mLNs and spleen at later time points, while earlier changes were primarily observed in pLNs. This suggests that rapamycin’s immunomodulatory effects are both time-dependent and site-specific.

**Figure 1.**
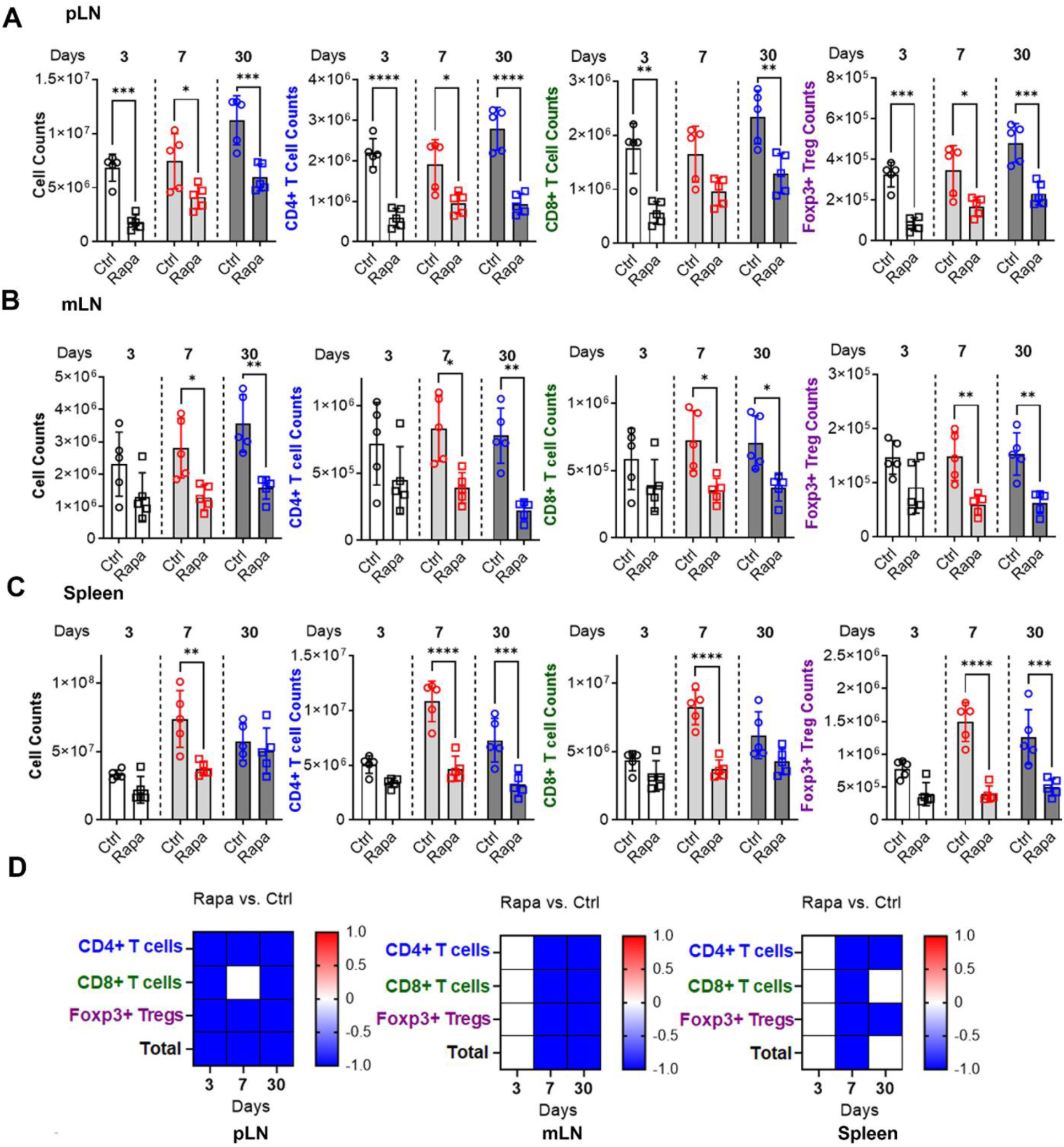
Rapamycin elicits significant changes in immune cell populations in LNs and spleen. Flow cytometry for the total number (CD45+ cells), CD4+ T cells, CD8+ T cells, and Foxp3+ Tregs (Foxp3+ CD4+) in **A)** pLN, **B)** mLN, and **C)** spleen after 3, 7, and 30 days of rapamycin treatment. **D)** Heatmap depicts changes in cell numbers relative to the control for pLN, mLN, and spleen after rapamycin treatment versus no drug control, red to represent “increased,” white to represent “unchanged,” and blue to represent “decreased.”. 5 mice/group. One-way ANOVA .* p < 0.05, ** p < 0.01, *** p < 0.001, **** p < 0.0001.

### Time-dependent impact of rapamycin on LN architecture and Treg distribution

The architecture and distribution of specific immune cells within LN microenvironments, including the CR and HEVs, are key in fostering immune tolerance and suppression (39). We conducted quantitative immunohistochemistry (IHC) of these LN domains to characterize rapamycin’s spatiotemporal effect.

Rapamycin rapidly induced early pro-inflammatory changes by decreasing the laminin α4:α5 ratio (La4:La5) in the pLN HEVs by day 3. Such reduction extended to the pLNs CR by day 7, and diminished to undetectable levels by day 30 **(Figure 2A, E, Supplemental Figure 2A, B)**, indicating an early pro-inflammatory impact on pLNs. In mLNs, rapamycin significantly increased La4:La5 on day 30, but no significant changes on days 3 and 7 **(Figure 2B, F, Supplemental Figure 2C)**, indicating a late pro-tolerogenic impact on mLN.

**Figure 2.**
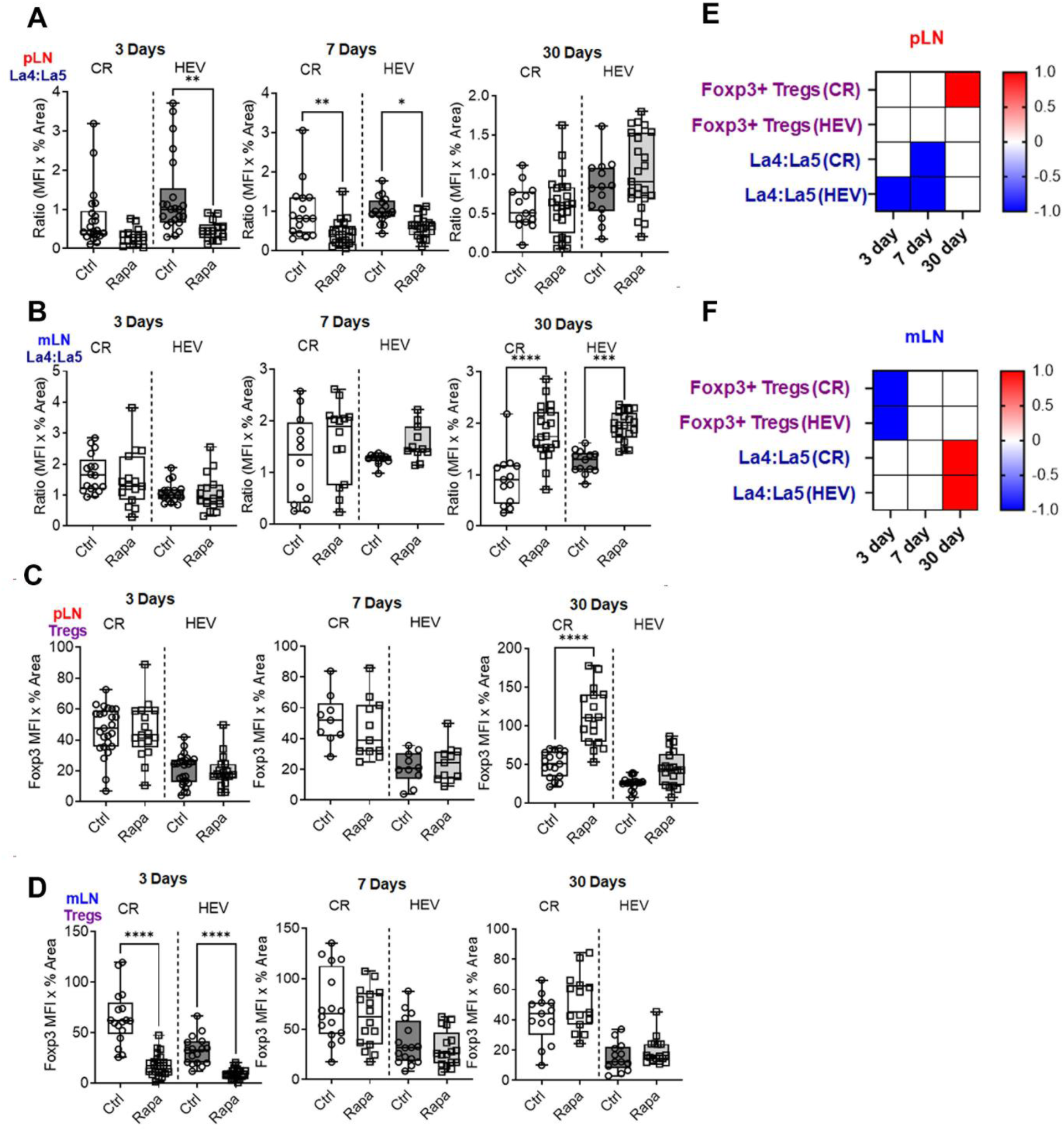
Effects of rapamycin on LN cell content, cell distribution, and structure. IHC of CR and HEV laminin α4:α5 (la4: la5) ratios on days 3, 7, and 30 of **A)** pLN and **B)** mLN. IHC of CR and HEV Foxp3+ Tregs on days 3, 7, and 30 of **C)** pLN and **D)** mLN. Heatmaps depict changes in expression with rapamycin relative to control in **E**) pLN and **F**) mLN. 3 mice/group. One-way ANOVA .* p < 0.05; ** p < 0.01, *** p < 0.001, **** p < 0.0001.

Rapamycin significantly increased Foxp3+ Tregs in pLNs by day 30, especially in the CR, while no alterations at early time points **(Figure 2C, E, Supplemental Figure 2D)**, indicating a late pro-tolerogenic effect on pLNs. In mLNs, rapamycin reduced Foxp3+ Tregs in the CR and around HEVs on day 3 but not at later time points **(Figure 2D, F, Supplemental Figure 2E)**, indicating an early pro-inflammatory impact on mLNs. Overall, both pLNs and mLNs exhibited early pro-inflammatory states, characterized by decreased La4:La5 or reduced Tregs. This was followed by a late pro-tolerogenic environment, marked by either increased La4:La5 or elevated Tregs. Notably, the data indicated that day 7 post-treatment marked a transition from pro-inflammatory to pro-tolerogenic regulation by rapamycin.

### Rapamycin increases laminin α5 and decreases La4:La5 in LNSCs

Given the impact of rapamycin on LN architecture via La4:La5, we next sought to determine whether stromal cell expression of laminin α4, α5, or both directly mediated these effects after 7 days treatment, given this time point marked the critical transition from pro-inflammation to pro-tolerance. Flow cytometry was used to quantify α4 and α5 levels in live CD45-LNSCs including: FRCs, BECs, and LECs. In pLNs, rapamycin upregulated laminin α5 in FRCs, LECs, and BECs, with little influence on laminin α4, leading to decreased laminin α4:α5 ratios **(Fig. 3A-G)**. In mLNs, treatment with rapamycin similarly decreased the laminin α4:α5 ratio in LNSCs by upregulating Laminin α5 expression **(Fig. 3H-M)**. Taken together, rapamycin modulates pro-inflammatory responses by selectively increasing laminin α5 expression in LNSCs and hence decreasing La4:La5 in LNSCs.

**Figure 3.**
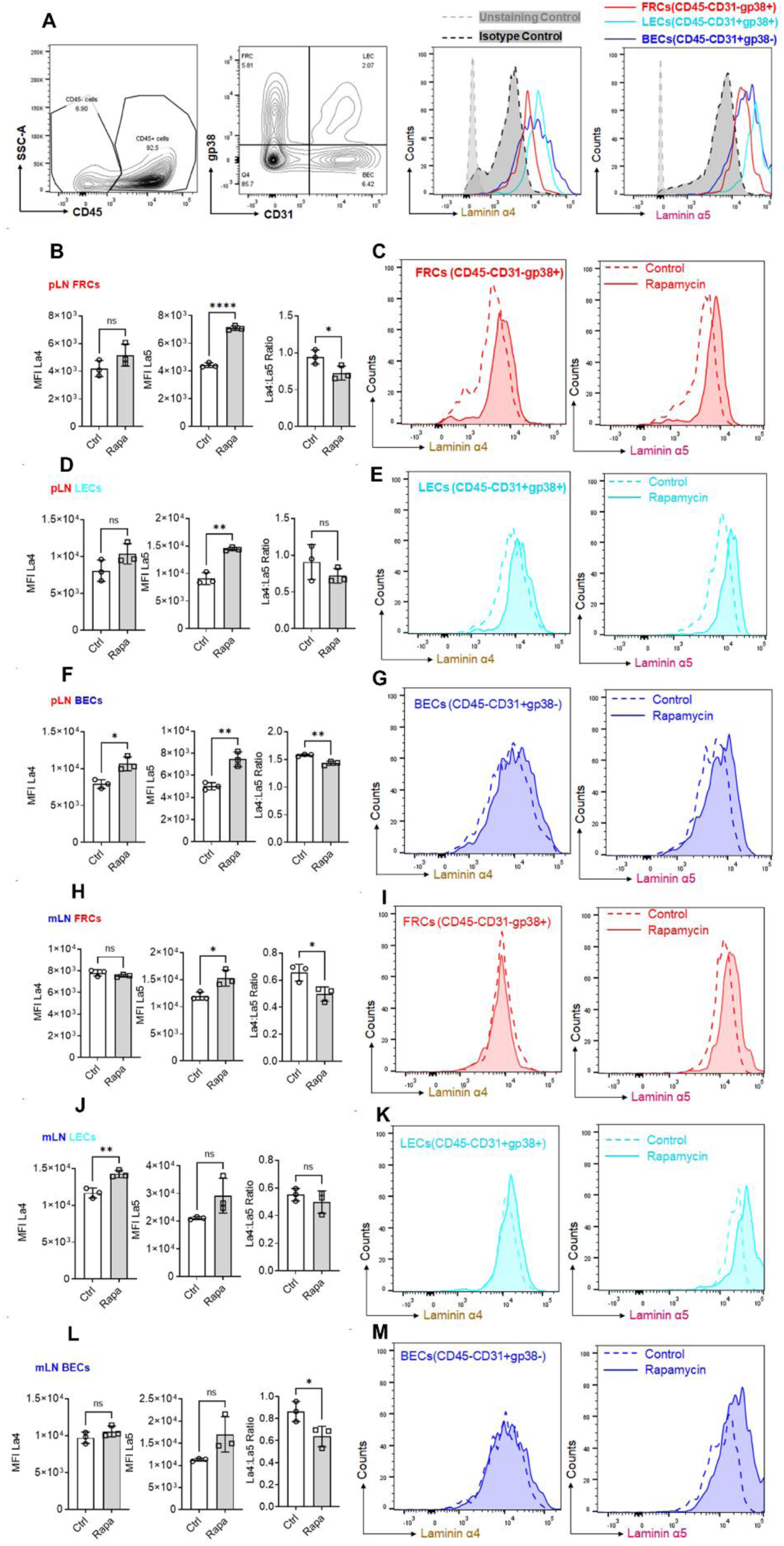
Rapamycin increases laminin α5 and decreases the laminin α4:α5 ratio in LNSCs. **A)** Flow gating of CD45-cells for FRCs (CD31-gp38+), BECs (CD31+gp38-), and LECs (CD31+gp38+), for laminin α4 (la4) and laminin α5(la5). MFI (Mean Fluorescence Intensity) and flow plots show laminin α4, laminin α5, and laminin α4 to laminin α5 ratios in: **B-G)** pLN: **B, C)** FRCs; **D, E)** LECs; and **F, G)** BECs and **H-M)** mLN: **H, I)** FRCs;**J, K)** LECs; and **L, M)** BECs. 3 mice/group. Unpaired t-test. * p < 0.05; ** p < 0.01, **** p < 0.0001.

### Laminin α5 is responsible for rapamycin-induced changes in LN architecture

Given the significant impact of rapamycin on pLNs La4:La5 on day 7 and the important role of FRCs in laminin expression, we further employed two laminin KO murine strains to assess if laminin α4 or α5 expression was mediated by FRCs. FRC-Laminin4-KO mice (Pdgfrb-Cre+/- × La4fl/fl) have the laminin α4 gene specifically deleted in FRCs. At baseline, these mice showed significantly decreased laminin α4 level in both pLNs and mLNs compared to WT mice **(Figure 4A, D)**. Administration of rapamycin did not affect the expressions of laminin α4 and laminin α5, and the La4:La5 in pLNs of the FRC-La4-KO mice **(Figure 4A, B, C)**. In mLNs, rapamycin significantly increased laminin α5 around HEV without affecting laminin α4 in FRC-Laminin4-KO, leading to a significantly reduced La4:La5 **(Figure 4E, F)**. These results indicate that even in the absence of FRC laminin α4, rapamycin can still upregulate laminin α5 expression, suggesting FRC-derived lama4 is not required to mediate the LN architectural change under the influence of rapamycin.

**Figure 4.**
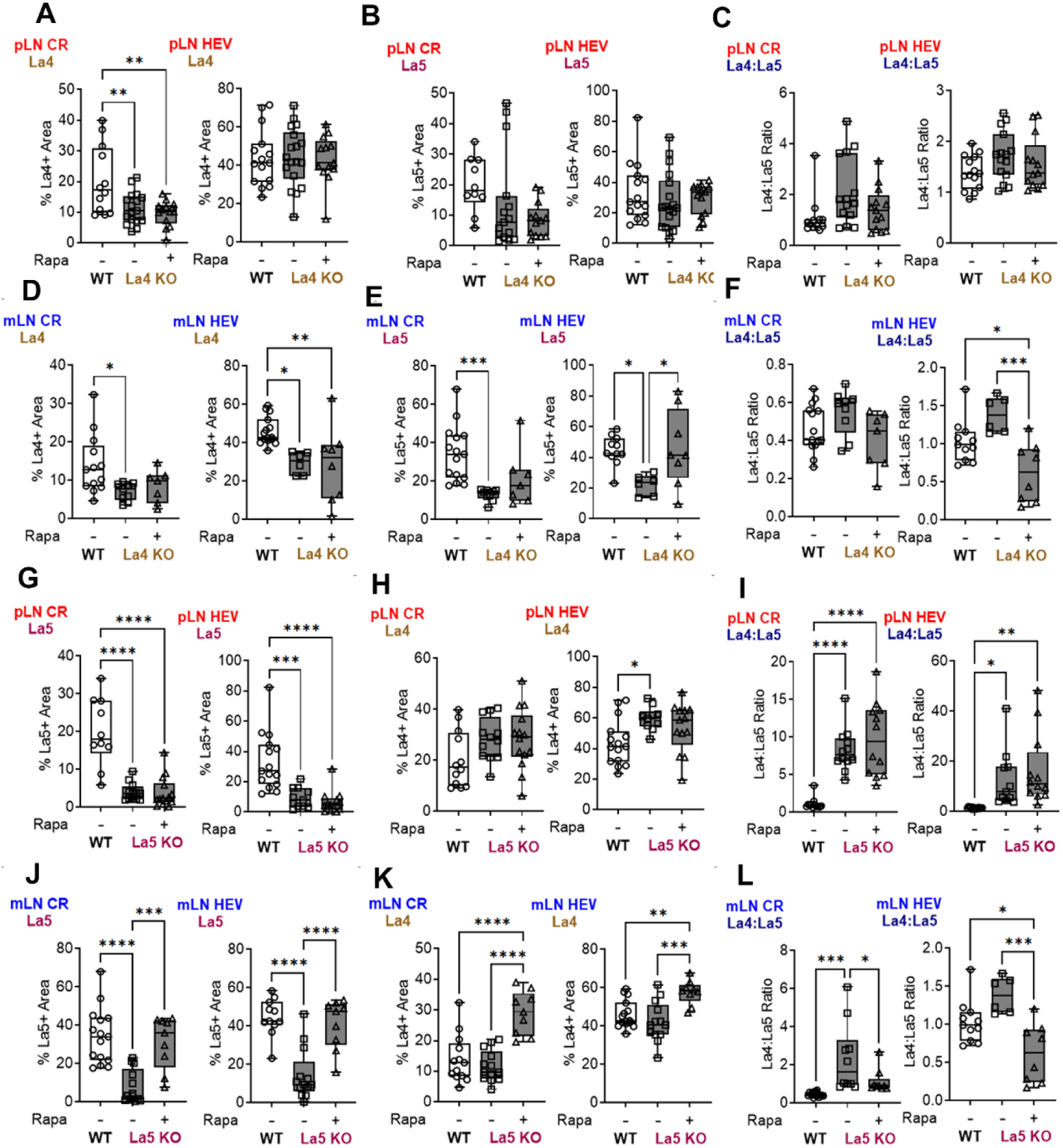
Rapamycin regulates laminin α5 in FRC-Lama4-KO and FRC-Lama5-KO mice. IHC showing A, **H)** laminin α4, **B, G)** laminin α5, **C, I)** laminin α4:α5 ratios in pLN, **D, K)** laminin α4, **E, J)** laminin α5, **F, L)** laminin α4:α5 ratios in mLN, for WT, untreated and rapamycin-treated FRC-Lama4-KO mice **(A-F)** and FRC-Lama5-KO mice **(G-L)** on day 3. 3-5 mice/group. One-way ANOVA. * p < 0.05, ** p < 0.01, *** p < 0.001, **** p < 0.0001.

FRC-Laminin5-KO mice (Pdgfrb-Cre+/- × La5fl/fl) have the laminin α5 gene specifically deleted in FRCs. At baseline, these mice showed significantly decreased laminin α5 in both pLN and mLN compared to WT mice **(Figure 4G, J)**. Administration of rapamycin did not affect the expression of laminin α4 or α5 in the pLN CR and around HEV, resulting in an unchanged La4:La5 in FRC-La5-KO mice **(Figure 4G, H, I)**. In the mLN, there was a differential increase in both laminin α4 and laminin α5 **(Figure 4J, K)**, leading to a decrease in the La4:La5 (**Figure 4L)**. Since the laminin α5 gene is selectively deleted in FRCs but not in other cells, the increased laminin α5 levels post-rapamycin treatment suggest upregulation of laminin α5 expression by non-FRC cell types, such as LECs and BECs. Flow cytometry results (**Figure 3**) support this conclusion. Overall, the results show that rapamycin differentially regulates laminin α4 and α5, altering LN architecture. This effect is not exclusively mediated by FRC-derived laminin, as non-FRC cells like LECs and BECs also play crucial roles in these rapamycin-induced changes.

### Time-dependent impact of rapamycin on gut microbiota composition and metabolic capacity

We next investigated rapamycin’s effect on the gut microbiome because of its critical role in interfacing with the immune system. Whole community metagenomic sequencing of intraluminal fecal contents was performed at a sequencing depth of 41.1±14.5 (mean±s.d.) million post-qc reads per sample (**Supplemental Table 2A**) and taxonomic composition was estimated using the comprehensive mouse microbiota genome catalog (44) (**Supplemental Table 2B**). After rapamycin treatment, no significant differences in gut microbiota diversity were observed at days 3 and 7 (**Figure 5A**). However, by day 30, there was a significant decrease in community diversity (P < 0.05) and a marked shift in microbial composition, characterized by increased *Bacteroides* (mainly *Muribaculaceae*) to *Firmicutes* (mainly *Lachnospiraceae*) relative abundance ratio, or B/F ratio (**Supplemental Figure 3A**). The B/F ratio on day 3 was 0.29±0.26 (mean±s.d.), which increased to 0.78±0.53 on day 7 and reached 1.19±1.13 by day 30, indicating a sustained impact of rapamycin on the structure and composition of the gut microbiota. Compared to the control group, no significant changes in taxonomic groups were observed on day 3. Sporadic alterations were noted on day 7 including *Bacteroidale* (*i.e.*, *Duncaniella sp.*, *Muricubaculum intestinale*) and *Firmicutes* (i.e., *Eubacterium*) (**Supplemental Figure 3B, Supplemental Table 2C**). By day 30, there was a marked increase in *Bacteroidales* (including *Muribaculaceae, Bacteroides, Actinobacteria*), and a decrease in Firmicutes (such as *Lachnospiraceae*, *Oscillibacter*, *Lawsonibacter*, *Eubacterium*) (**Supplemental Figure 3C**). These taxonomic groups drive the observed changes in composition and structure of the gut microbiota at different times and between groups (**Figure 5B**). These effects, starting at the intermediate time point of day 7 and becoming most pronounced by day 30, highlight a sustained and potent influence of rapamycin on the microbiome, and correspond to the same time frame for changes in LN immune architecture and content.

**Figure 5.**
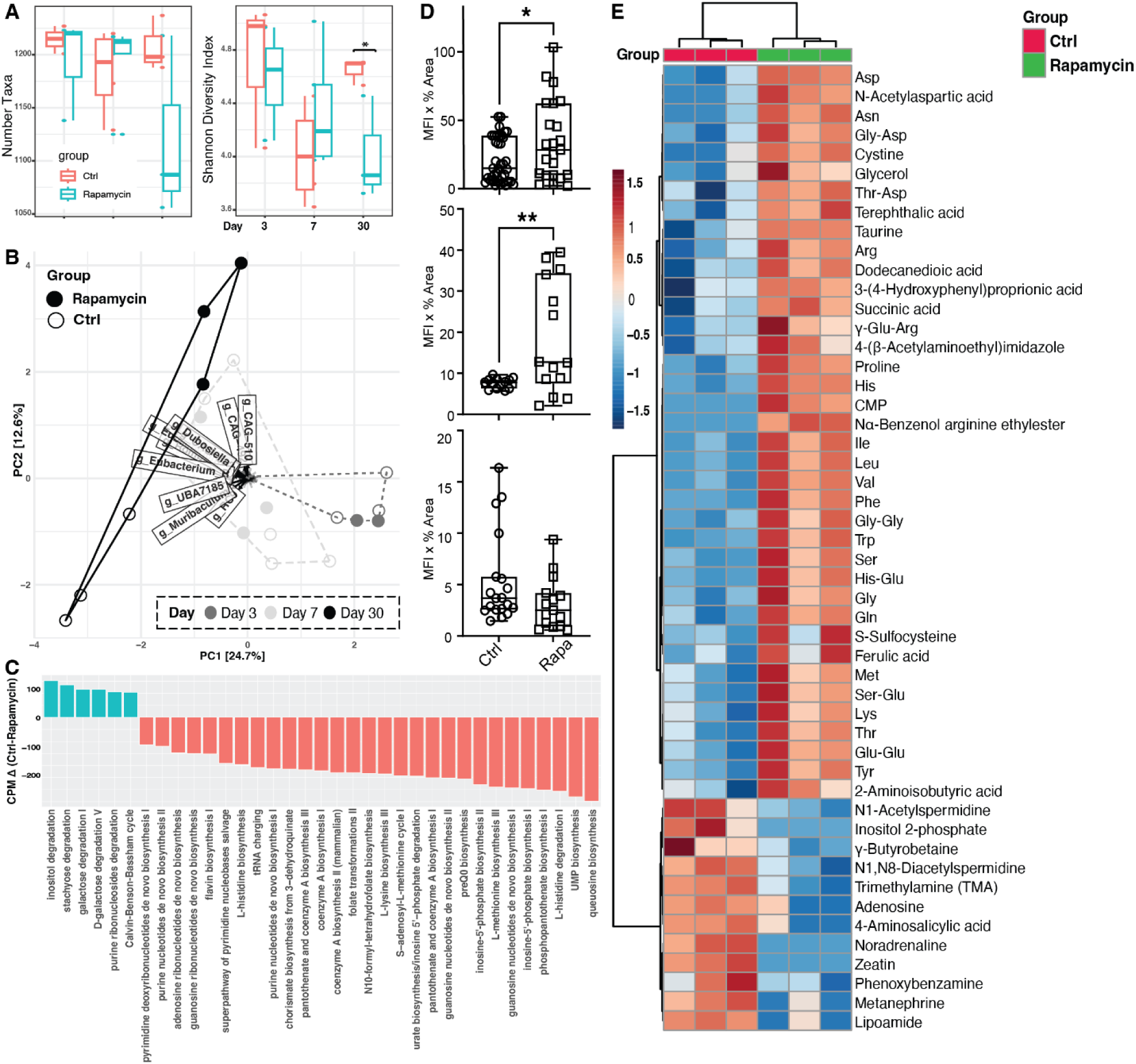
Rapamycin alters gut microbiome and metabolic potentials. Gut microbiome characterization of rapamycin treated and no treatment control for **A)** within-community diversity by total number of taxa and Shannon diversity index; **B)** community beta-diversity with PCA; and **C)** gut microbiome functional pathway abundance in copies per million (CPM) difference between control and rapamycin groups. The height of the stacked bar represents CPM of associated MetaCyc pathways contributed by different taxa in control (green color) or rapamycin (red color). Intestinal IHC of **D)** Foxp3+ Tregs after 3 days, 7 days, and 30 days of rapamycin treatment. **E)** Hierarchical clustering heatmap of metabolites of rapamycin and control groups. Specimens were collected after rapamycin treatment for 7 days, compared to no drug control. Top 50 features shown. Color bar indicates the scaled z-score of each feature.

Functional pathway characterization revealed a significant increase in nucleotide biosynthesis and reduction in glycolysis and multiple sugar degradation metabolic pathways after 30 days of rapamycin treatment (**Figure 5C**). These changes were not significant after 3 or 7 days of treatment. Species-resolved functional pathway analysis revealed that the *Muribaculaceae* family (Bacteroides) was enriched after long-term rapamycin treatment, compared to control, and harbored pathways involved in nucleotide biosynthesis (**Supplemental Figure 4A**). On the other hand, *B. thetaiotaomicron* and a variety of Clostridiales taxa (Firmicutes) that were relatively depleted after long-term rapamycin treatment harbored functional pathways in amino acid biosynthesis (*i.e.*, ornithine), branched and aromatic amino acid biosynthesis (horismate pathway), glycolysis and energy processing (**Supplemental Figure 4B**). These findings revealed a distinct shift in gut microbiome’s composition and function following rapamycin treatment, emphasizing rapamycin’s impact beyond immuunosupression and a potential mechanism for its diverse therapeutic effect.

### Rapamycin temporally shifts intestinal immune responses

To elucidate the reciprocal interactions between the gut microbiome and host under rapamycin treatment, we analyzed the intestinal transcriptome. Days 7 and 30 were chosen as they corresponded to the major gut microbiota alterations following rapamycin treatment. Differentally expressed genes (DEGs) were identified by comparing the rapamycin group to the no-treatment control. A total of 69 and 234 upregulated DEGs were observed at days 7 and 30, respectively; and a total of 168 and 84 down-regulated DEGs were observed at days 7 and 30, respectively (**Supplemental Table 3A,B**). At day 7, 70.9% of DEGs were down-regulated, while at day 30, 73.6% of DEGs were up-regulated. On the other hand, intestinal Foxp3+ Treg expression was strongest at days 3 and 7, but attenuated by day 30 (**Figure 5A-C**). The enriched upregulated immune pathways at both days 7 and 30 include B cell regulation, activation, proliferation, antigen binding, and immunoglobulin mediated immune responses (**Supplemental Figure 5A**). On Day 7, there was unique enrichment in cellular responses to interferon-λ, -α, and -β, as well as cytokine-mediated signaling pathways. At day 30, unique enrichment was observed in MHC class II protein complex binding, antigen processing and presentation, mucosal immune responses, and tissue-specific immune responses. Other immune pathways demonstrated a substantially stronger effect in most functional categories by day 30, such as immunoglobulin receptor binding, production, and circulation, positive regulation of lymphocyte activation, and phagocytosis (**Supplemental Figure 5B**). These results collectively indicate a temporal shift from suppression to activation in both intestinal gene expression and the immune environment following rapamycin treatment.

### Rapamycin reprograms amino acid metabolism in gut lumen

Given the significant alterations in the composition and functional makeup of the gut microbiome, we next sought to assess whether this translated to functional changes in metabolism through through the gut luminal metabolome. Intraluminal stool was assessed using capillary electrophoresis-mass spectrometry (CE/MS) (45–47). The 7-day time point was used as it represents the transitional phase between early and late alloimmune responses in both LNs and the intestine. A no-drug treatment group served as a control to provide a baseline for comparison. Luminal metabolites (n=264) were exhaustively annotated by PubChem (48), Kyoto Encyclopedia of Genes and Genomes (KEGG) (49), and Human Metabolome Database (HMDB) (50) (**Supplemental Table 4A**). According to the KEGG BRITE hierarchical classification system, the most prevalent class of luminal metabolites was from amino acid metabolism, comprising 42.7% of all annotated metabolites (**Supplemental Table 4B**). These metabolites belonged to pathways of arginine and proline, histidine, tyrosine, and tryptophan metabolism. Other prevalent classes included carbohydrates (11.5%), cofactors and vitamins (10.4%), nucleotides (10.4%), lipids (6.3%), other amino acids (6.3%), and xenobiotic metabolism (5.2%). Distinct gut metabolic profiles were observed after rapamycin treatment, with amino acids such as Asn, Phe, Arg, and Leu and metabolic derivatives differentially abundant in rapamycin treatment group (**Figure 5D, Supplemental Figure 6A-B**). These results suggest rapamycin treatment either increased amino acid biosynthesis and/or reduced catabolism.

### Rapamycin induces a rapid pro-inflammatory response and gut microbiome shift during allogeneic stimulation

We next employed a mouse model with allogeneic stimulation (allo) to characterize the effect of rapamycin on transplant-related alloimmune responses. Mice were injected with fully allogeneic splenocytes (10^7^ cells intravenously) followed by rapamycin treatment on day 0. A concise 3-day post-treatment window was used given the rapid and sensitized immune responses. Compared to allo alone and no treatment control, allo alone leads to a decrease in La4:La5 in pLNs, consistent with our previous findings (51). When allostimulation was combined with rapamycin treatment, there was an increase in both laminin α4 and α5 compared to allogeneic splenocytes alone. However, the increase in laminin α5 was more significant than in laminin α4, resulting in decreased laminin α4:α5 ratios in both the pLNs and mLNs **(Figure 6A-F)**. This pattern mirrored findings from WT and KO mice treated with rapamycin without allostimulation. Further, allogeneic splenocytes increased pLNs CR and HEV Tregs without changes in mLNs Tregs **(Figure 6G, H)**. After 3 days of treatment with rapamycin plus allostimulation, compared to allostimulation alone, there was no alteration in the distribution of Tregs in pLNs, but Tregs were decreased in the mLNs CR and HEV **(Figure 6G-I),** indicating a pro-inflammatory state. These data demonstrated that rapamycin also fostered an early pro-inflammatory LN environment in the context of alloantigen-induced immune responses through altering La4:La5 and Treg distribution.

**Figure 6.**
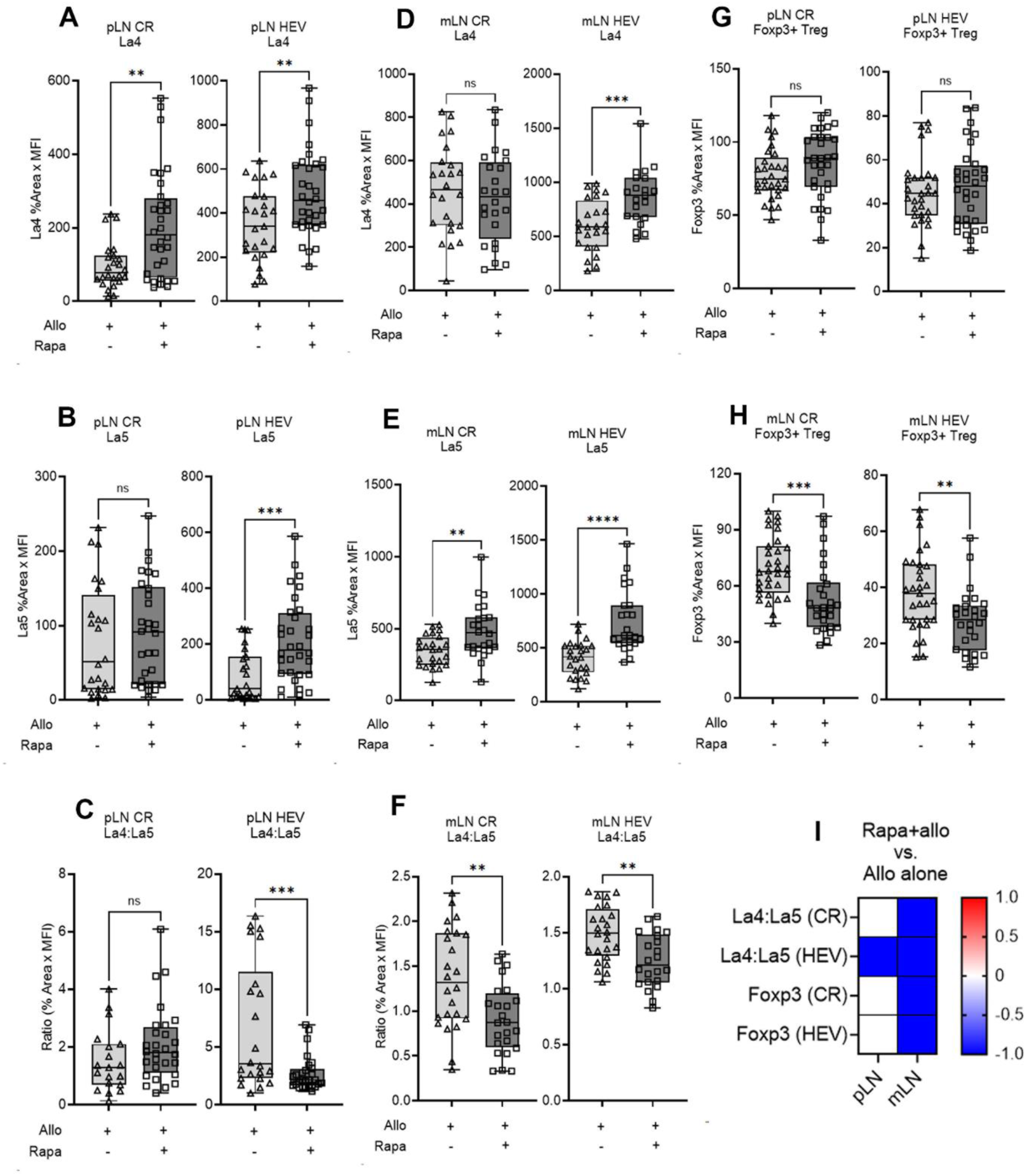
Rapamycin triggers a rapid inflammatory immune response in mice immunized with allogeneic splenocytes. IHC for pLN **A)** laminin α4, **B)** laminin α5, **C)** laminin α4:α5 ratios. IHC for mLN **D)** laminin α4, **E)** laminin α5, **F)** laminin α4:α5 ratios. CR and HEV Foxp3+ Tregs in **G)** pLN and **H)** mLN. **I)** Heatmap changes in marker expression comparing allostimulation plus rapamycin to allostimulation only in pLN and mLN. 3 mice/group. One-way ANOVA. * p < 0.05, ** p < 0.01, *** p < 0.001, **** p < 0.0001. allo - allogeneic splenocytes.

The combination of allostimulation and rapamycin treatment for three days led to significant changes in the gut microbiome, characterized by increased microbial diversity and altered community composition and structure. In contrast, allostimulation alone showed no notable differences compared to the untreated control group (**Figure 7A, B, Supplemental Table 5**). While the total number of microbial taxa remained unchanged, the Shannon diversity index significantly increased in the rapamycin plus allostimulation group (P < 0.05). This suggests that rapamycin, in the context of alloantigen-induced immune responses, promotes a more even distribution of microbial species without altering the overall number of distinct taxa. A marked shift in microbiome composition was observed, accompanied by this increase in microbial diversity. The B/F ratio decreased substantially in the rapamycin plus allogeneic stimulation group (0.12±0.10) compared to untreated controls (0.45±0.22) or allostimulation alone (1.16±0.67), indicating a significant restructuring of the microbial community (**Figure 7B, Supplemental Figure 7A**). Allostimulation alone increased the relative abundance of potentially pro-inflammatory *Muribaculaceae* (i.e., *Duncaniella* and *Paramuribaculum*) within the Bacteroides phylum. However, the combination of rapamycin and allostimulation led to a higher abundance of Firmicutes, including *Lachnospiraceae, Butyricicoccaceae,* and CAG-274 (**Figure 7C,D, Supplemental Figure 7B-D**).

**Figure 7.**
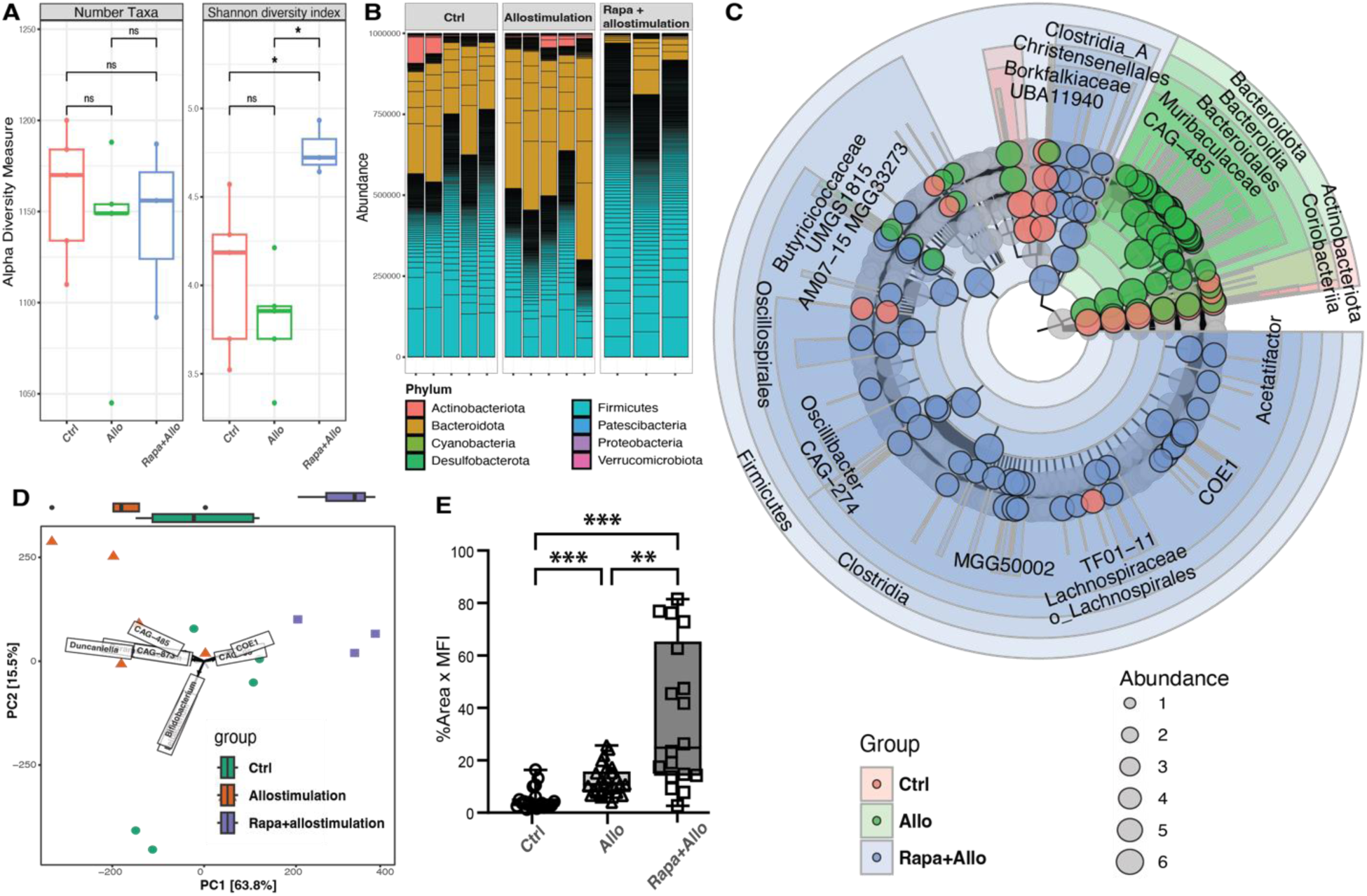
Effects of rapamycin and alloimmunity on gut microbiome and intestinal Foxp3+ Treg. Gut microbiome characterization of no treatment control, allostimulation, and rapamycin combined with allostimulation in **A)** diversity index using observed number of taxa and Shannon diversity index; **B)** taxonomic composition on phylum level; and **C)** cladogram of differentially abundant taxonomic groups using LDA effect size (LEfSe). Each filled circle represents one biomarker. The diameter of a circle is proportional to the phylotype relative abundance scaled at log 10. **D)** PCA plot with taxa loadings labeled. Length of each taxa loading vector indicates its contribution to each PCA axis shown. The univariable group distribution plotted as side panel. **E)** IHC of Foxp3 Tregs in intestine assessed after 3 days of rapamycin treatment in allostimulated mice.

These results demonstrated the rapid, phylogenetic-aware impact of rapamycin in an alloantigen-stimulated environment where multiple taxa within the same phylogenetic group swiftly shifted in a unified direction. While similar to observations in naïve mice under rapamycin treatment, the specific taxonomic groups affected differed. Furthermore, allostimulation significantly increased intestinal Foxp3+ Tregs, an effect further enhanced by rapamycin treatment (**Figure 7E**). This pattern mirrors changes seen in naïve mice under rapamycin treatment with increased intestinal Tregs. Collectively, these findings highlight rapamycin’s context-dependent influence on the intestinal microenvironment, affecting both intestinal Treg populations and gut microbiome during allogeneic immune challenges.

## DISCUSSION

This study was designed to capture early, intermediate, and late effects of rapamycin, to reveal time-dependent and site-specific immunomodulatory effects (**Figure 8**). The results demonstrated the dynamics of rapamycin’s influence on immune responses and the gut microbiome, from the intestine to mesenteric and peripheral LNs, showing coordinated, multifaceted impact across different anatomical sites and time points. Locally, in the intestine, rapamycin affected gut microbiota, intestinal immune cell responses, and luminal metabolic activities, transitioning from an early pro-tolerogenic to a late pro-inflammatory state. Regionally, in the mLNs, rapamycin modulated immune responses and stromal cell function. The mLNs act as critical intermediaries, reflecting gut-originating changes that impact regional immune responses, demonstrated by dynamic changes in laminin α4:α5 ratios and Treg distribution. Systemically, in the pLNs, rapamycin significantly altered immune cell populations and LN architecture, mirroring the regional transitions from pro-inflammatory to pro-tolerogenic states. This indicated that rapamycin’s effects, initiated locally in the gut, rapidly propagated to regional and distant lymphoid tissues and impacted overall immune homeostasis. The overlapping effects of rapamycin treatment over time and space illustrate a compartmentalized yet coordinated immune response, emphasizing the need to consider both targeted and broad impacts when evaluating immunosuppressive therapies.

**Figure 8.**
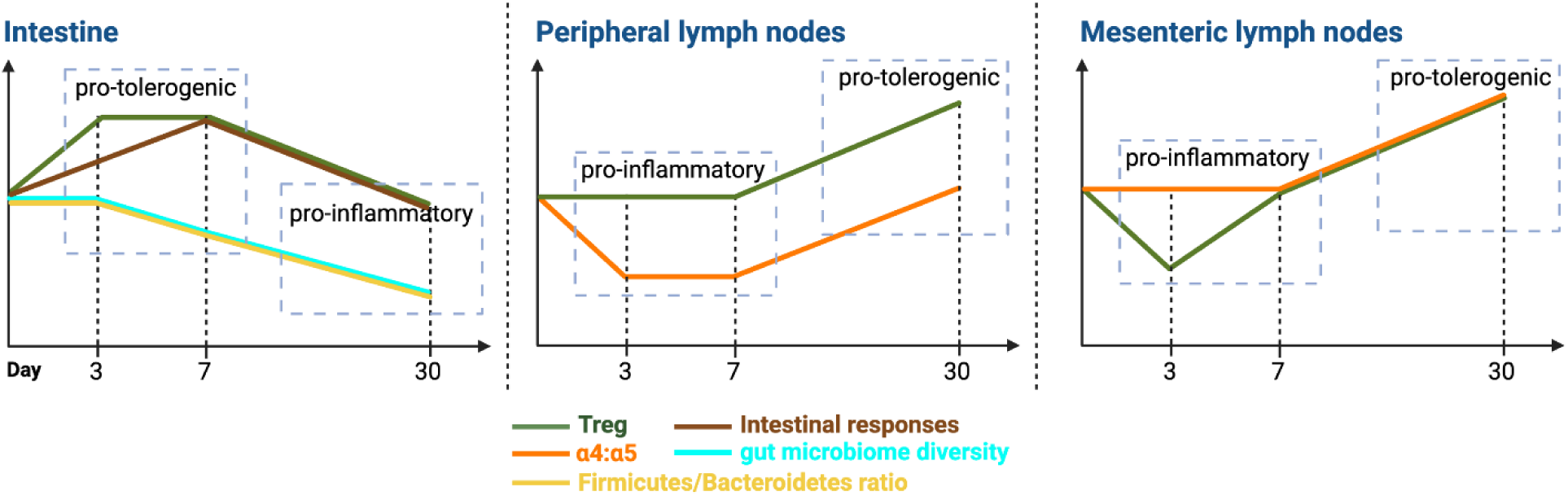
Pattern of changes in intestine, pLN, and mLN. The results of longitudinal changes summarized to demonstrate the temporal changes over time. Illustration generated in Biorender.com

Rapamycin has well-documented immunosuppressive effects on both innate and adaptive immune responses through the inhibition of mTOR signaling (3, 4). However, mechanisms underlying these effects remains incompletely understood. In this study, we demonstrated that rapamycin facilitated a significant transition between pro-inflammatory and pro-tolerogenic states by selectively increasing laminin α5 expression in LNSCs. Our previous research found that laminin α5 inhibits Treg cells (51). These findings collectively suggest a more complex mechanism of rapamycin effects that involves a reduction in Tregs and a concurrent LN restructuring. This finding was validated across models, including naïve, laminin α4 KO, laminin α5 KO, and allogeneic splenocyte-stimulated mice. Rapamycin’s influence on LNSCs likely represents a key mechanism of its action, as LNSCs critically regulate lymphocytes positioning and interaction with antigen-presenting cells via chemokines, cytokines, and stromal fibers (52–54). By altering LNSC function, rapamycin may reshape the spatial organization and dynamics of immune cell interactions, thereby modulating overall immune responses.

Rapamycin’s regulation of LN architecture extended to include LECs and BECs. This broad impact on the LNSC network further emphasizes rapamycins’ role in modulating the LN function and structure. LECs play a critical role in lymph drainage and immune cell trafficking, while BECs regulate the entry of circulating lymphocytes into the LN. By modulating these cell types, rapamycin could be altering the trafficking patterns and retention of various immune cell subsets within the LN, thereby influencing the initiation and progression of immune responses. Overall, our results highlight its multifaceted mechanisms that act not just on immune cells directly but also on the structural and functional elements of lymphoid organs. This enhanced understanding may inform the analysis of immune mechanisms during the development of new immunosuppressive drugs where precise immune modulation is crucial.

The effects of rapamycin on the gut microbiome were incremental and depended on the duration of exposure. Over time, a temporal shift in intestinal responses and the immune environment was observed, transitioning from suppression to activation following prolonged drug use. One key change was the reprogramming of amino acid metabolism in the gut lumen. This change was at least partially due to alterations in the microbiome, specifically a few keystone species that may play a major role in driving these metabolic modifications. Since amino acids are essential for mTORC1 activation, their increased bioavailability in the gut lumen may represent a potential mechanism by which rapamycin influences alloimmune responses. However, the causal relationship between rapamycin treatment, gut microbiome alterations, and immune responses remains unclear. It is yet to be determined whether the observed changes in microbiota composition and function are a direct result of rapamycin’s effect or an indirect consequence of rapamycin-induced changes in immune and intestinal responses. Future experiments involving immune-deficient mice and fecal microbiota transfer from rapamycin-treated subjects following varied treatment intervals are warranted.

The accumulation of amino acids in the gut lumen during rapamycin-induced immunosuppression is a complex phenomenon attributable to multiple interrelated factors. First, rapamycin’s inhibition of mTORC1 plays a critical role in regulating protein synthesis (8, 55), as mTORC1 normally promotes translation initiation and protein synthesis through the activation of downstream targets like p70S6 Kinase I (S6K1) and elF4E Binding Protein (4EBP). This inhibition leads to an accumulation of amino acids that would otherwise be utilized in protein synthesis. Second, mTORC1 inhibition induces autophagy, facilitating the breakdown of intracellular components including proteins by lysosomal degradation, potentially increasing the pool of free amino acids for cell survival (56). Third, rapamycin’s suppression of immune cell proliferation and activity further contributes to amino acid accumulation by decreasing metabolic demand. In particular, we observed the reduction in metabolic products like N1-acetylspermidine and the increase in arginine bioavailability suggesting reduced arginine catabolism due to rapamycin treatment. Fourth, rapamycin’s disruption of the gut microbiome alters the availability of gut and circulatory amino acids, influencing host nutrient homeostasis and physiology (57). Fifth, rapamycin also directly affects amino acid metabolism in the intestine by modulating synthesis, utilization, and transport processes (58, 59). Further investigation into the metabolic effects of rapamycin could inform targeted nutritional and supportive interventions to optimize rapamycin’s therapeutic efficacy.

## MATERIALS & METHODS

### Study approval

All procedures involving mice were performed in accordance with the guidelines and regulations set by the Office of Animal Welfare Assurance of the University of Maryland School of Medicine under the approved Institutional Animal Care and Use Committees (IACUC) protocol nos. 05150001, 0318002, 1220001, and AUP-00000397.

### Mice experiments

Female C57BL/6 mice between 8 and 14 weeks of age were purchased from The Jackson Laboratory (Bar Harbor, ME, USA) and maintained at the University of Maryland School of Medicine Veterinary Resources animal facility. We only used female mice for the current set of experiments where we worked with smaller n-values, to ensure a high degree of homogeneity within our study groups. The Pdgfrb-Cre+/– x La4fl/fl (42) and Pdgfrb-Cre+/– x La5fl/fl (38) conditional knockout (KO) mice were previously developed in our laboratory. To simulate allogeneic stimulation, mice received 10^7 fully allogeneic BALB/c splenocytes intravenously (i.v.) on day 0 and rapamycin. Rapamycin (USP grade, MilliporeSigma) was reconstituted in DMSO (USP grade, MilliporeSigma) at 25 mg/mL. DSMO stock was diluted in sterile phosphate buffered saline (PBS) to 0.5 mg/ml for intraperitoneal (i.p.) injection at 5 mg/kg/d, following protocols from prior investigations (43, 60). All mice were cohoused and handled together during arrival in the animal facility and for immunosuppressant administration so that the various treatment groups were all exposed to each other. On the day of harvest, the mice were euthanized by CO_2_ narcosis, intraluminal stool samples collected for metabolomic analyses, cardiac puncture utilized for blood collection, and mesenteric and peripheral (axillary, inguinal, and brachial) LNs and small intestine harvested for immunological assays.

### Flow cytometry

LNs and spleens were disaggregated and passed through 70-μm nylon mesh screens (Thermo Fisher Scientific, Waltham, MA) to obtain single-cell suspensions. The digestion protocols for FRCs, LECs, and BECs analysis in pLN and mLN were followed as outlined previously (61). The cell suspensions were treated with anti-CD16/32 (clone 93, eBioscience) to block Fc receptors and stained for 15 minutes at 4°C with antibodies targeting surface molecules **(Supplemental Table 1)** and washed 2 times with buffer (PBS with 0.5% w/v bovine serum albumin). Cells were permeabilized using Foxp3/Transcription Factor Staining Buffer Set (eBioscience, San Diego, CA) according to manufacturer’s protocol, washed with buffer, and subsequently stained at 4°C with antibodies for intracellular molecules. Samples were analyzed with an LSR Fortessa Cell Analyzer (BD Biosciences), and data were analyzed using FlowJo software version 10.8.1 (BD Biosciences). Single color controls (cells stained with single surface marker antibody) and unstained controls were used for flow channel compensation.

### Immunohistochemistry

mLN and pLN, and segments of intestine between the duodenum and jejunum were separately excised and immediately submerged in OCT compound (Sakura Finetek, Torrance, CA). Sections (6 μm for LNs, 10 μm for intestine) were cut in triplicate using a Microm HM 550 cryostat (Thermo Fisher Scientific) and fixed in cold 1:1 acetone:methanol for 5 minutes, and washed in PBS. Primary antibodies (**Supplemental Table 1**) were diluted between 1:100 and 1:200 in PBST (PBS + 0.03% Triton-X-100 + 0.5% BSA) and incubated for 1 hour in a humidified chamber. Sections were then washed with PBS, and secondary antibodies (**Supplemental Table 1**) were applied at a 1:400 dilution in PBST for 1 hour. Slides were washed in PBS for 5 minutes, coverslipped, and imaged using an Accu-Scope EXC-500 fluorescent microscope (Nikon, Tokyo, Japan) and the images analyzed with Volocity software (PerkinElmer, Waltham, MA). 3-6 mLNs and 3-6 pLNs and 3 pieces of intestine, 3-6 sections/tissue sample, and 10-20 fields/tissue sample were analyzed. Mean of mean fluorescence intensity (MFI) was calculated within demarcated HEVs and CR regions of mLN and pLN as well as of whole intestinal images. Percent area was calculated by dividing the sum area of demarcated regions with marker fluorescence greater than a given threshold by total area analyzed. Treatment groups were compared using quantitation of MFI multiplied by percent area (%Area x MFI) to express both area and intensity of cell and stromal fiber markers.

### Stool specimen collection, DNA extraction, and metagenomic sequencing

Stool pellets were collected on days 0, 3 and then at weekly intervals and at termination of the experiment for longitudinal characterization. Intraluminal stools were collected from colon at harvest. They were stored immediately in DNA/RNA Shields (Zymo Research, Irvine, CA, USA) and archived at −80°C. DNA extraction was described previously (28, 62). In brief, 0.15-0.25 grams of stool samples were extracted using the Quick-DNA Fecal/Soil Microbe kit (Zymo Research, Irvine, CA, USA). Negative extraction controls were included to ensure that no exogenous DNA contaminated the samples. Metagenomic sequencing libraries were constructed using the Nextera XT Flex Kit (Illumina, San Diego, CA, USA), according to the manufacturer’s recommendations. Libraries were then pooled together in equimolar proportions and sequenced on a single Illumina NovaSeq 6000 S2 flow cell at Maryland Genomics at the University of Maryland School of Medicine.

### Gut microbiome analyses

Metagenomic sequence reads mapping to Genome Reference Consortium Mouse Build 39 of strain C57BL/6J (GRCm39) (63) were removed using BMTagger v3.101 (64). Sequence read pairs were removed when one or both the read pairs matched the genome reference. The Illumina adapter was trimmed and quality assessment was performed using default parameters in fastp (v.0.21.0) (65). The taxonomic composition of the microbiomes was established using Kraken2 (v.2020.12) (66) and Braken (v. 2.5.0) (67) using the comprehensive mouse microbiota genome catalog (44). Phyloseq R package (v1.38.0) (68) was used to generate the barplot and diversity index. Linear discriminant analysis (LDA) effect size (LEfSe) analysis (69) was used to identify fecal phylotypes that could explain the differences. The α value for the non-parametric factorial Kruskal-Wallis (KW) sum-rank test was set at 0.05 and the threshold for the logarithmic LDA model (70) score for discriminative features was set at 2.0. An all-against-all BLAST search was performed in the multiclass analysis. Phylogram representing the taxonomic hierarchical structure of the identified phylotype biomarkers via pairwise comparisons between groups, graph generated using R package yingtools2 (71). Taxonomic ordination graphs were created with the microViz (v0.12.4) (72). The metagenomic dataset was mapped to the protein database UniRef90 (73) to ensure the comprehensive coverage in functional annotation, and was then summarized using HUMAnN3 (Human Microbiome Project Unified Metabolic Analysis Network) (v0.11.2) (74) to determine the presence, absence, and abundance of metabolic pathways in a microbial community. MetaCyc pathway definitions and MinPath were used to identify a parsimonious set of pathways summarized in HUMAnN3 in the microbial community. The total pathway abundance was further stratified by contributing species in HUMAnN3. MaAsLin2 (75) was used to identify the association of the pathways with groups.

### RNA isolation, transcriptome sequencing and analyses of the intestinal tissues

The colon tissues of the mice were dissected according to their anatomic features and stored immediately in RNAlater solution (QIAGEN) in RNAse-free 1.7-ml tubes (Denville Scientific, Holliston, MA) at −80°C to stabilize and protect the integrity of RNA. For each sample, total RNA was extracted from ∼1 cm of ileum. Prior to the extraction, 500 µl of ice-cold RNase free PBS was added to the sample. To remove the RNAlater, the mixture was centrifuged at 8,000x*g* for 10 min and the resulting pellet resuspended in 500 µl ice-cold RNase-free PBS with 10 µl of β-mercaptoethanol. A tissue suspension was obtained by bead beating procedure using the FastPrep lysing matrix B protocol (MP Biomedicals, Solon, OH) to homogenized tissues. RNA was extracted from the resulting suspension using TRIzol Reagent (Invitrogen, Carlsbad, CA) following the manufacturer’s recommendations and followed by protein cleanup using Phasemaker tubes (Invitrogen) and precipitation of total nucleic acids using isopropanol. RNA was resuspended in DEPC-treated DNAase/RNAase-free water. Residual DNA was purged from total RNA extract by treating once with TURBO DNase (Ambion, Austin, TX, Cat. No. AM1907) according to the manufacturer’s protocol. DNA removal was confirmed via PCR assay using 16S rRNA primer 27 F (5′-AGAGTTTGATCCTGGCTCAG-3′) and 534 R (5′- CATTACCGCGGCTGCTGG-3′). The quality of extracted RNA was verified using the Agilent 2100 Expert Bioanalyzer using the RNA 1000 Nano kit (Agilent Technologies, Santa Clara, CA). Ribosomal RNA depletion and library construction were performed using the RiboZero Plus kit and TruSeq stranded mRNA library preparation kit (Illumina) according to the manufacturer’s recommendations. Libraries were then pooled together in equimolar proportions and sequenced on a single Illumina NovaSeq 6000 S2 flow cell at the Genomic Resource Center (Institute for Genome Sciences, University of Maryland School of Medicine) using the 150 bp paired-end protocol. An average of 87-130 million reads were obtained for each sample. The quality of Fastq files was evaluated by using FastQC (76). Reads were aligned to the mouse genome (Mus_musculus.GRCm39) using HiSat (version HISAT2-2.0.4,) (77) and the number of reads that aligned to the coding regions was determined using HTSeq (78). Significant differential expression was assessed using DEseq2 (79) with an FDR value ≤ 0.05 and Fold-of-change (FC)>2. The over-representative analysis was done by importing Differentially Expressed Genes (DEGs) against GO ontologies using the enrichGO function of clusterProfile Bioconductor package (80). Only the ontology terms with q <0.05 were used for plotting.

### Metabolome analyses

Metabolome of intraluminal stool samples collected from ileum was measured using capillary electrophoresis-mass spectrometry (CE/MS) to obtain a comprehensive quantitative survey of metabolites (Human Metabolome Technologies, Boston, MA, USA). ∼10-30 mg of stool was weighed at the time of collection using a company-provided vial and archived at −80°C at the IGS until shipped to the HMT on dry ice. QC procedures included standards, sample blanks and internal controls that were evenly spaced among the samples analyzed. Compound identification was performed using a CE/MS library of >1,600 annotated molecules. Log base 10 transformation was applied on data to reduce the influence of measurement noise (81). Metabolites were annotated using PubChem (48), KEGG (49), and HMDB (50) annotation frameworks that leverage cataloged chemical compounds, known metabolic characterization, and functional hierarchy (i.e., reaction, modules, pathways). The sparse PLS-DA (sPLS-DA) algorithm implemented using mixOmics (vers. 6.18.1) was employed to analyze the large dimensional datasets that have more variables (metabolites) than samples (p >> n) to produce robust and easy-to-interpret models (82). The “sparseness” of the model was adjusted by the number of components in the model and the number of variables within each component based on the classification error rate with respect to the number of selected variables. Tuning was performed one component at a time, and the optimal number of variables to select was calculated. The volcano plot combines results from FC analysis to show significantly increased metabolites after 7-day tacrolimus treatment. A metabolite is shown if FC is >2 and the p-value is <0.05 based on 2-sample t-tests. Original metabolite measurements without normalization were used in the FC analysis.

### Data availability

The data that support the findings of this study are openly available; metagenome sequences were submitted to GenBank under BioProject PRJNA809764 (https://www.ncbi.nlm.nih.gov/bioproject/PRJNA809764) with the SRA study ID SRPXXX. The R codes, including each step and parameters, were deposited in github at https://github.com/igsbma/XXX. scRNASeq data were deposited in the NCBI Gene Expression Omnibus database (GEO XXX). Qualitative heat maps were generated (GraphPad prism) to express changes in marker expression level relative to control using 1 to represent “increased,” 0 to represent “unchanged,” and −1 to represent “decreased.”

### Statistics

The experiments were performed in three separate trials, with each trial containing at least three samples. Datasets were analyzed using GraphPad Prism 10.2.3 (San Diego, CA, USA) with statistical significance defined as *P* < 0.05. For comparisons of fluorescent markers (including laminin α4:α5 ratios), immune cell population distribution by flow cytometry, and immunohistochemistry markers, T tests or one-way ANOVA were used to test for significance.

## Authors’ contributions

L. W., B. M., and J. S. B. designed the experiments. L. W., A. K., S. J. G., R. L., D. K., L. L., V. S., W. P., and W. S. conducted and analyzed the *in vitro* and *in vivo* experiments. H. W. L. performed nucleic acid extraction and sequencing library preparation. B. M. and Y. S. performed the bioinformatics analyses. B. M., L. W., M. F., S. J. G, V. R. M., and J. S. B. wrote the manuscript. B. M. and J. S. B. are co-corresponding authors for this manuscript.

## ACKNOWLEDGEMENTS

This study was supported by 1R01HL148672 (JSB, BM), 1R01AI114496 (JSB), U01AI170050 (JSB, BM, VM), training grant T32AI95190-10 (SJG), and the University of Maryland Baltimore, Institute for Clinical & Translational Research (ICTR) (1UL1TR003098).

## Abbreviations

APC: Antigen Presenting Cell
ATC: Allogeneic splenocytes
BEC: Blood Endothelial Cell
CR: Cortical Ridge
Ctrl: Control
DC: Dendritic Cell
FRC: Fibroblastic Reticular Cell
HEV: High Endothelial Venule
IHC: Immunohistochemistry
KO: Knockout
La4: Laminin α4
La5: Laminin α5
LEC: Lymphatic Endothelial Cell
LN: Lymph Node
LNSC: LN Stromal Cell
mLN: mesenteric Lymph Nodes
mTORC1: mTOR Complex 1
mTORC2: mTOR Complex 2
pLN: peripheral Lymph Nodes Rapa Rapamycin
Treg: Regulatory T Cell

## Supplemental Figures

**Supplemental Figure 1.**
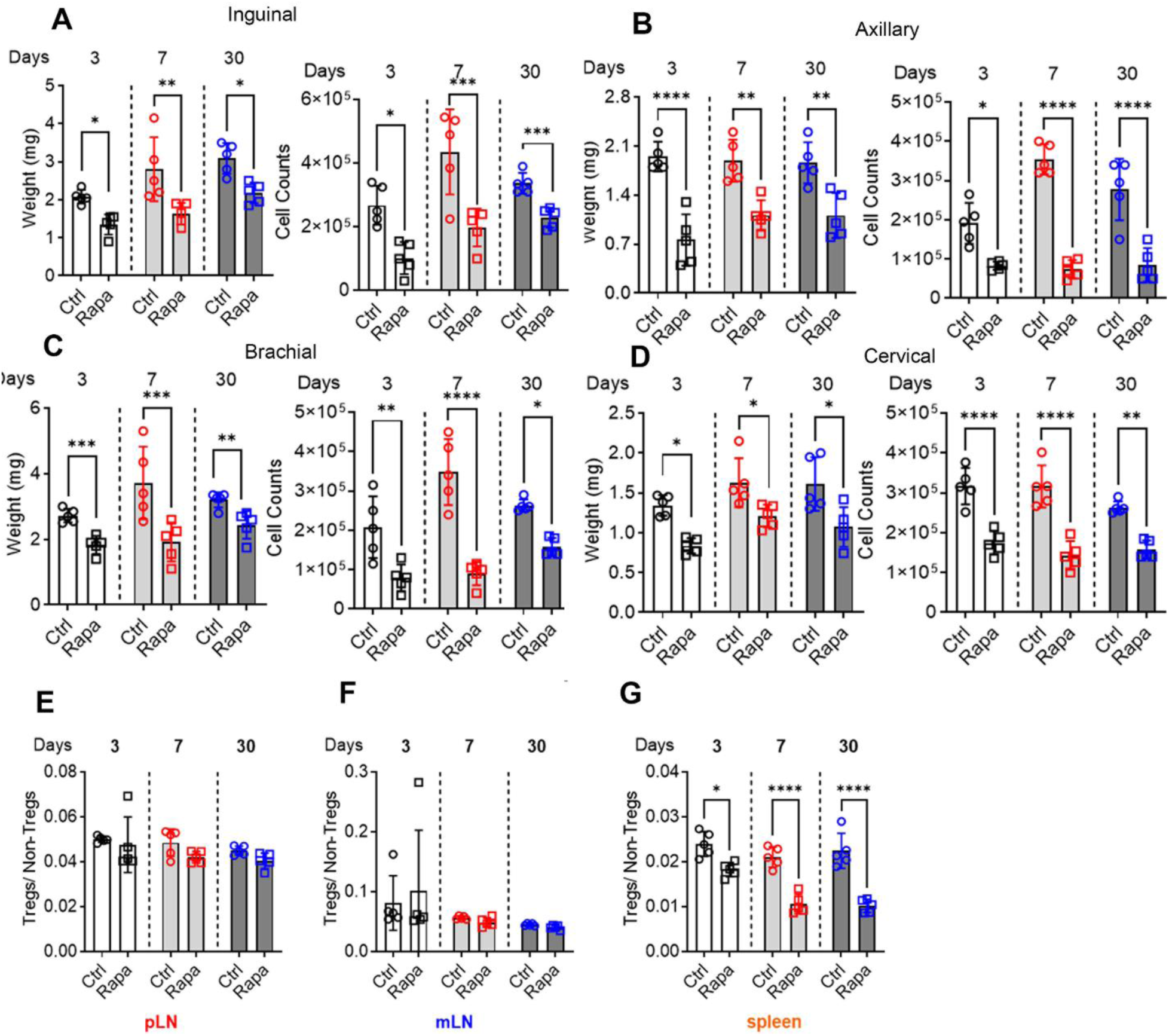
Rapamycin reduces cell counts in pLN. Weight and cell number for **A**) inguinal, **B)** axillary, **C)** brachial, and **D)** cervical LNs. Ratios of Tregs to non-Tregs in the **E)** pLN, **F)** mLN, and **G)** spleen. 5 mice/group. One-way ANOVA. * p < 0.05, ** p < 0.01, *** p < 0.001, **** p < 0.0001.

**Supplemental Figure 2.**
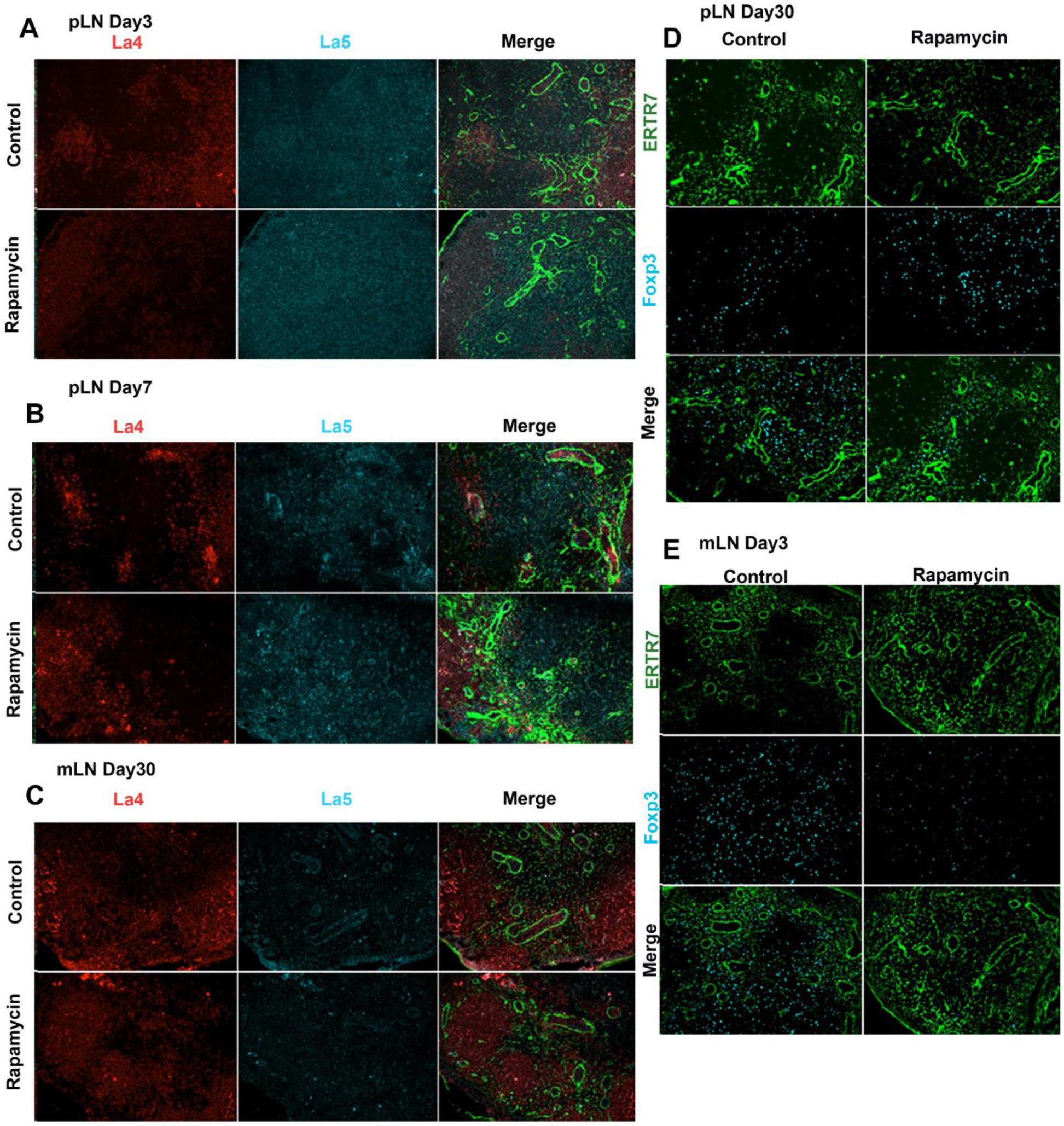
Effects of rapamycin on LN cell content, cell distribution, and structure. IHC images of Laminin α4, Laminin α5 for **A**) pLN on day 3, **B)** pLN on day 7, and **C)** mLN on day 30. ERTR7, Foxp3 for **D**) pLN on day 30 and **E**) mLN on day 3.

**Supplemental Figure 3.**
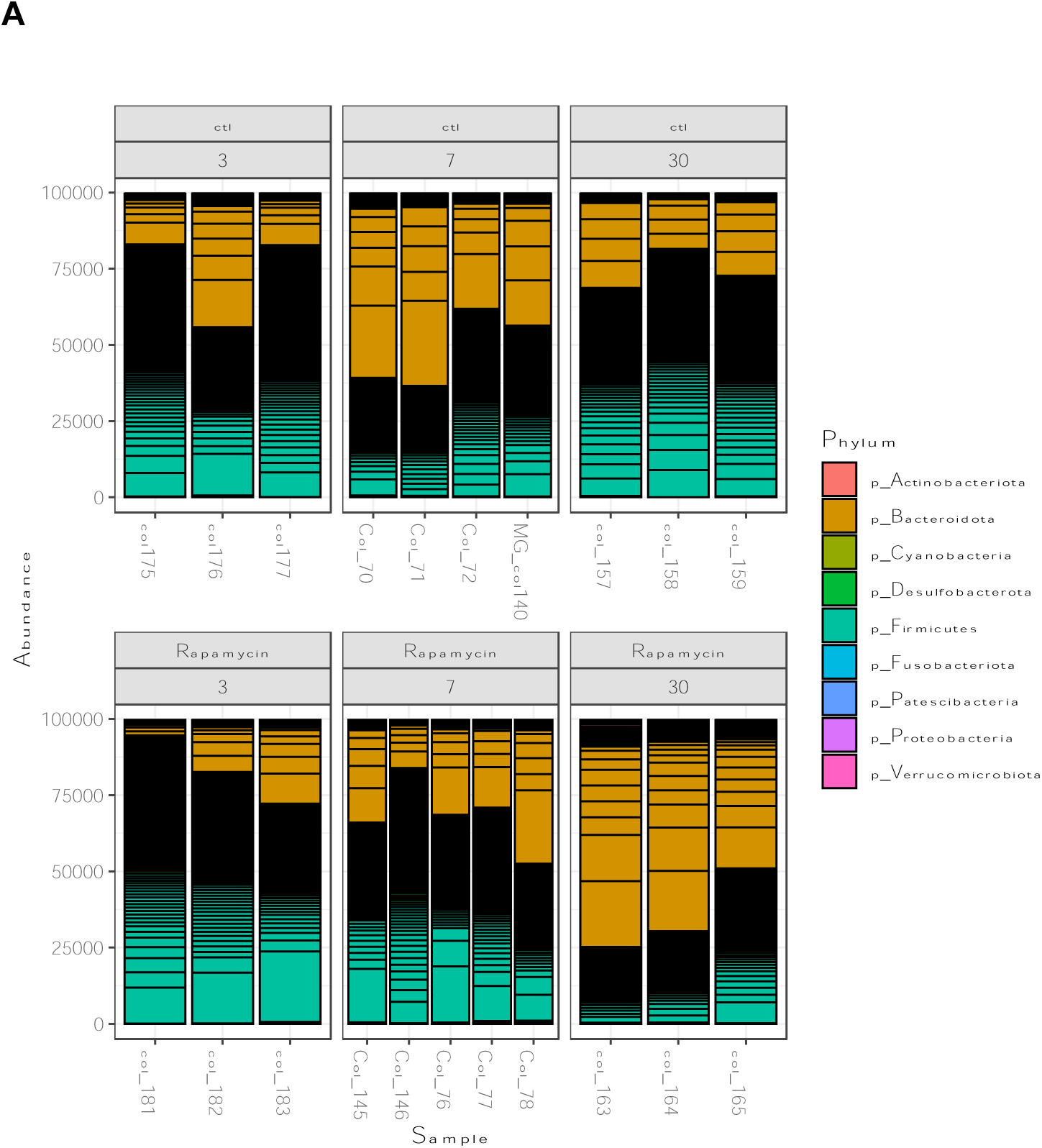
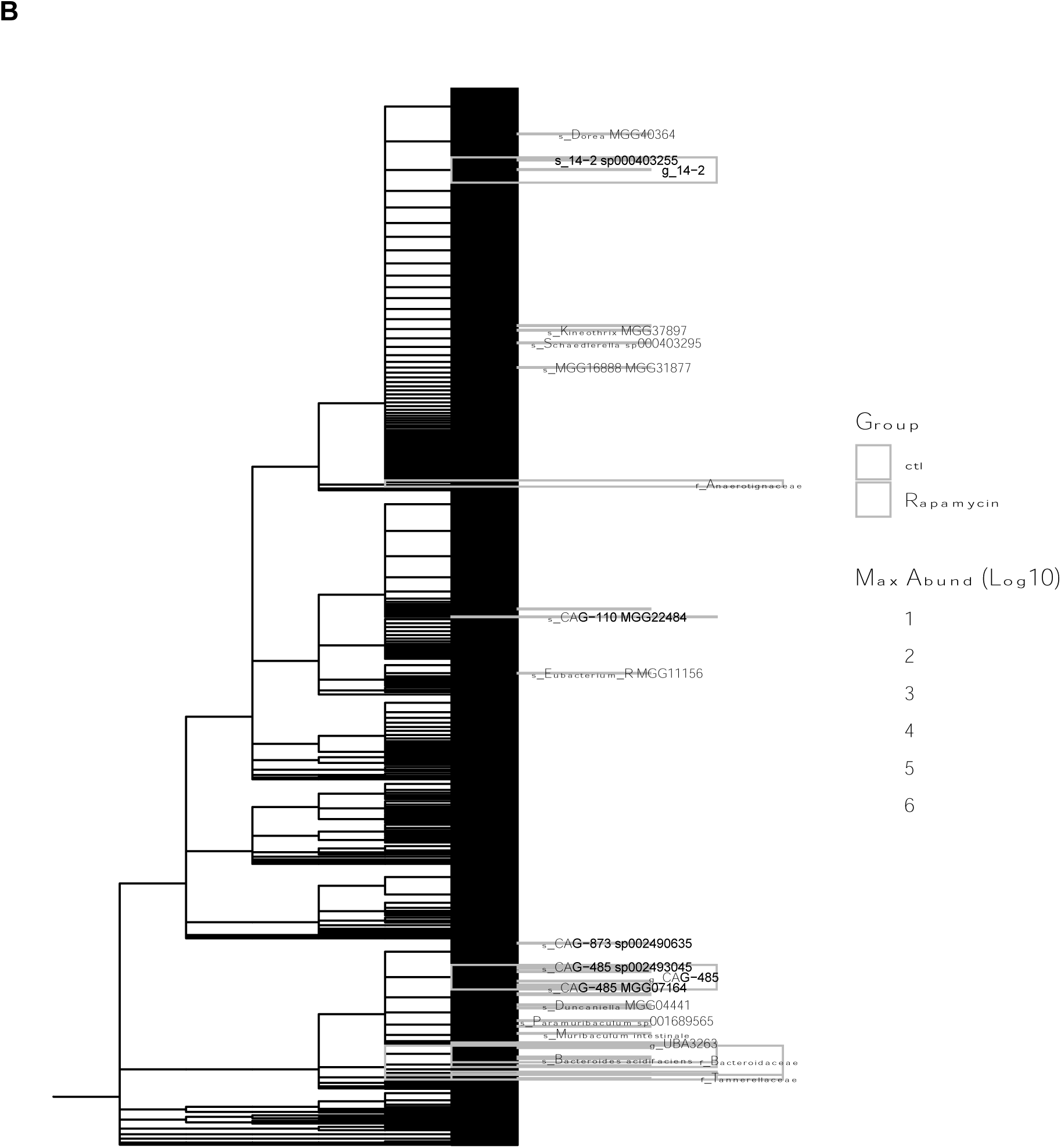
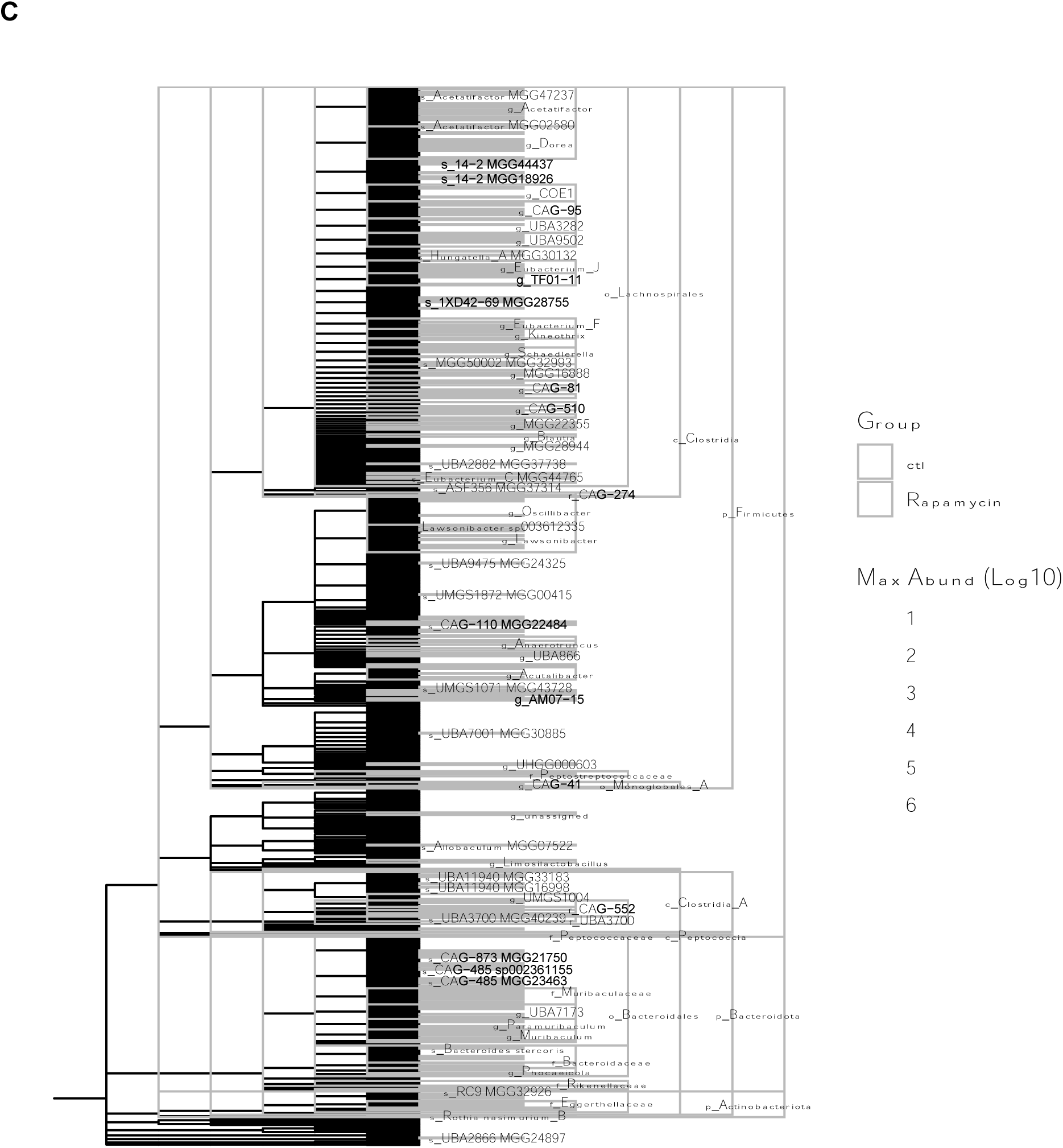
Rapamycin increases Firmicute of gut microbiome. **A)** Taxonomic composition of gut microbiome at phylum level after rapamycin treatment for 3, 7 and 30 days compared to no treatment control. Logarithmic linear discriminant analysis (LDA) effect size (LEfSe) (Segata et al., 2011) of the identified phylotype biomarkers of **B)** 7-day treatment; and **c**) 30-day treatment.

**Supplemental Figure 4. Rapamycin effect on the functional pathways of gut microbiome and contributing species.** Significantly enriched functional pathways in **A)** rapamycin group and **B)** no treatment control group, and responsible taxonomic groups in gut microbiome. Functional pathway composition and abundance generated using UniRef90 (Kanehisa et al., 2012) and HUMAnN2 (Franzosa et al., 2018) following MetaCyc pathway definitions (Caspi et al., 2014) and MinPath (Ye and Doak, 2009), and resulting pathways were stratified by contributing species. Figures are available to download at https://doi.org/10.6084/m9.figshare.26343112.

**Supplemental Figure 5.**
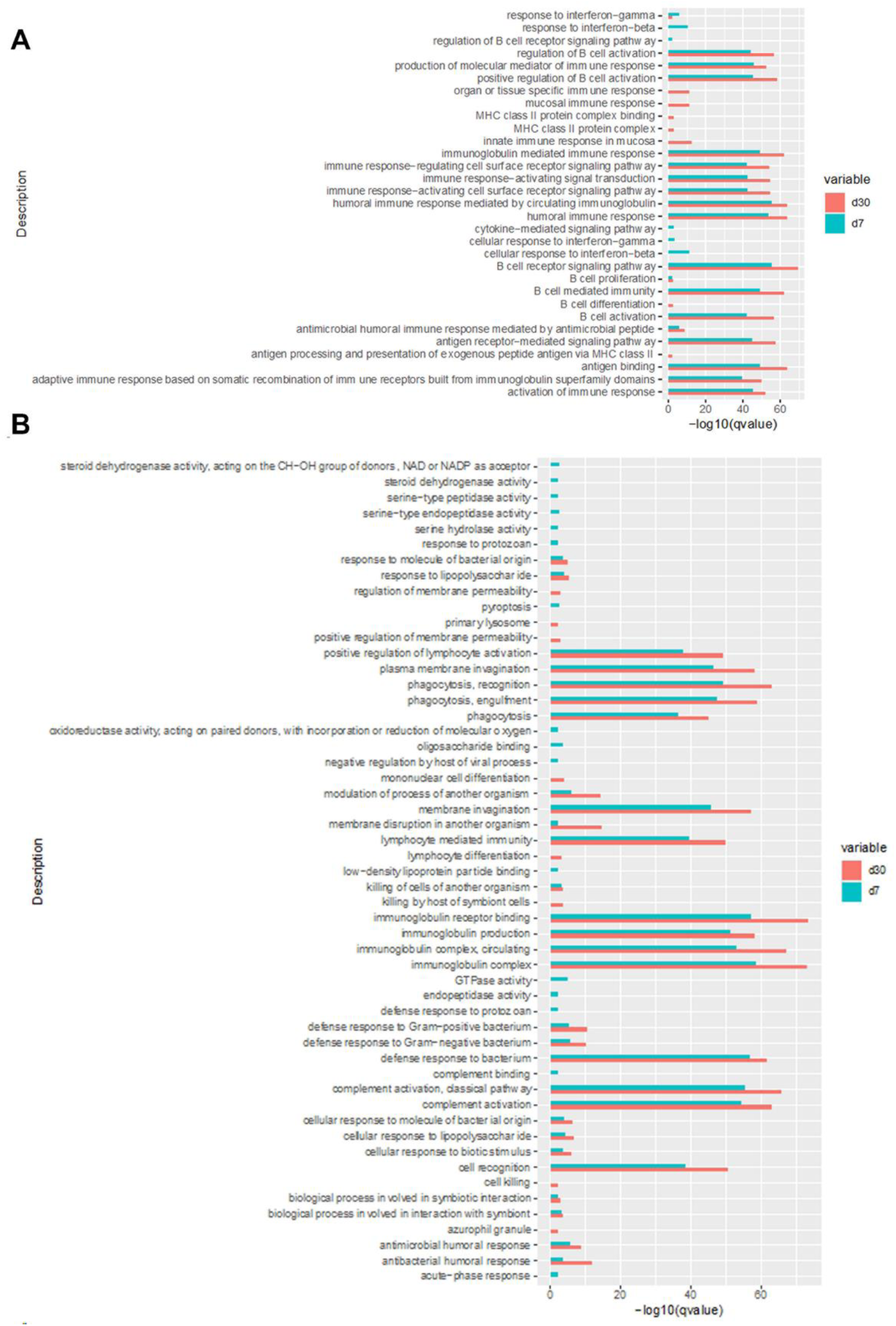
Rapamycin elicited non-immune intestinal responses. Top enriched **A**) immune-related and **B**) non-immune functional pathways of differential expressed genes of small intestinal tissues in rapamycin treated group after 7 and 30 days compared to no treatment control. Significant differential expression assessed using DEseq2 (Anders and Huber, 2010), the over-representative analysis performed by importing Differentially Expressed Genes (DEGs) against GO ontologies using the enrichGO function (G. Yu et al., 2012).

**Supplemental Figure 6.**
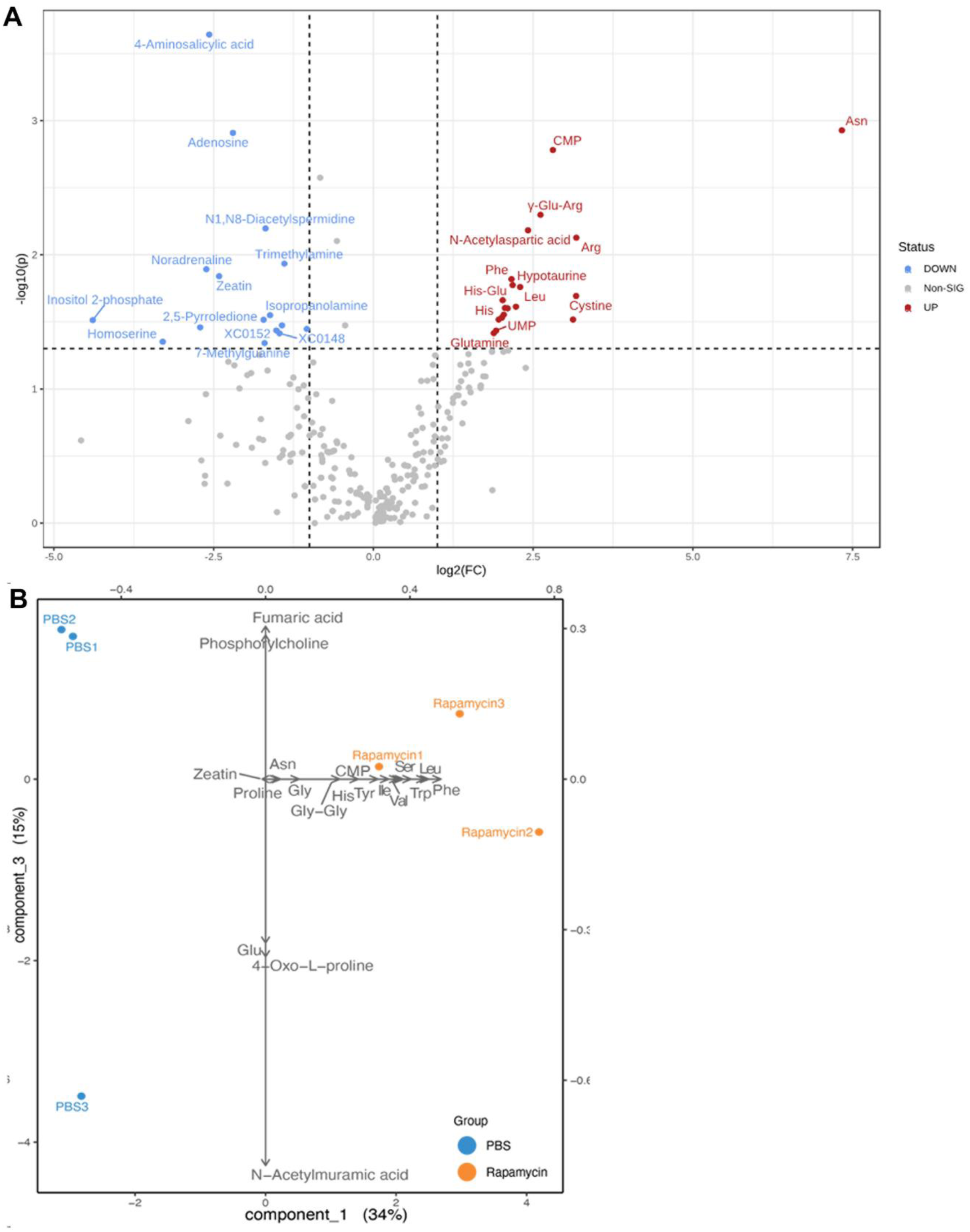
Differentially abundant intraluminal metabolites under rapamycin treatment. **A**) Volcano plot combines results from fold change (FC) analysis to show significantly increased metabolites after rapamycin treatment for 7 days compared to no treatment control. Metabolites shown if FC is >2 and p value <0.05 based on 2-sample t-tests. Original metabolite measurement without normalization used in FC analysis. **B)** Biplot of intraluminal stool metabolome. Loading vectors and principal components labeled.

**Supplemental Figure 7.**
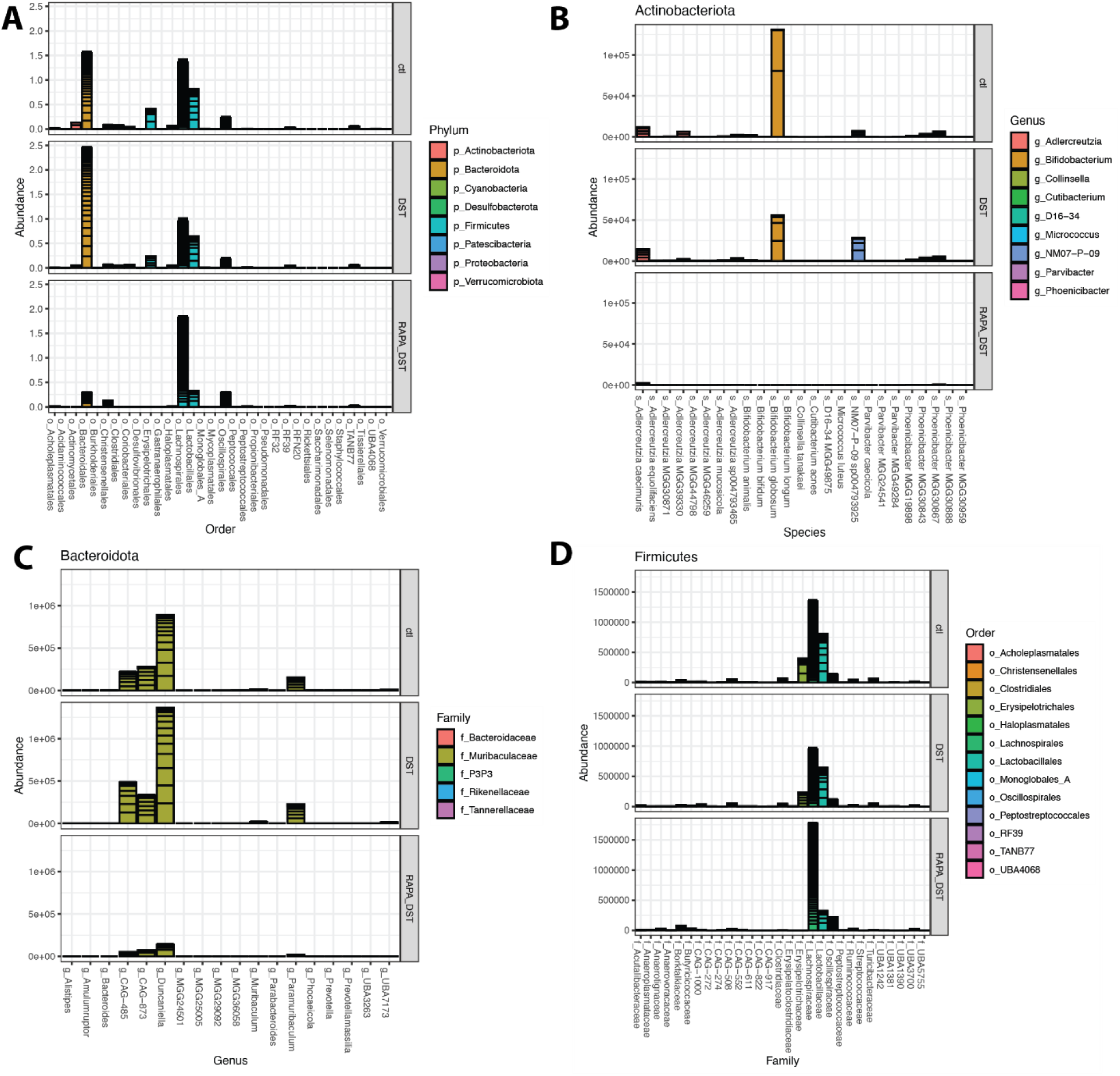
Rapamycin influence on gut microbiome of alloantigen sensitized mice. Comparison of relative abundance of bacterial groups in gut microbiota of alloimmune sensitized mice without and with rapamycin. Cumulative relative abundance of **A**) all taxonomic groups at order level, **B**) Actinobacteriota, **C**) Bacteroidota, and **D**) Firmicutes

## Supplemental Tables

**Supplemental Table 1.** List of Antibodies.

| Target Molecule | Clone | Catalog No. |
| --- | --- | --- |
| Anti-Hamster IgG Cy3 | Polyclonal | Jackson ImmunoResearch; 127-165-160 |
| Anti-Mouse IgG-AF647 | Polyclonal | Jackson ImmunoResearch; 715-605-151 |
| Anti-Rabbit AF488 | Polyclonal | Jackson ImmunoResearch; 111-545-003 |
| Anti-Rabbit Cy5 | Polyclonal | Jackson ImmunoResearch; 111-175-003 |
| Anti-Rabbit DL405 | Polyclonal | Jackson ImmunoResearch; 711-476-152 |
| Anti-Rabbit IgG-AF488 | Polyclonal | Jackson ImmunoResearch; 711-545-152 |
| Anti-Rabbit IgG-AF594 | Polyclonal | Jackson ImmunoResearch; 111-585-003 |
| Anti-Rabbit IgG-AF647 | Polyclonal | Jackson ImmunoResearch; 711-606-152 |
| Anti-Rat IgG AF594 | Polyclonal | Jackson ImmunoResearch; 112-586-143 |
| Anti-rat IgG AF647 | Polyclonal | Jackson ImmunoResearch; 712-606-153 |
| CD4 | GK1.5 | Biolegend; 100401 |
| CD45 | 30-F11 | Biolegend; 103132 |
| CD31 | 390 | Biolegend; 102427 |
| CD8 | 53-6.7 | Biolegend; 100701 |
| ER-TR7 | ER-TR7 | Novus; NB100-64932 |
| ER-TR7 | Polyclonal | Santa Cruz; SC-73355 |
| Foxp3 | NRRF-30 | eBioSci; 14477180 |
| Foxp3 | FJK-16s | eBioSci; 12-5773-82 |
| Gp38 | 8.1.1 | Biolegend; 127412 |
| Laminin α4 | 775830 | R&D; MAB3837 |
| Laminin α5 | Polyclonal | Novus Biol; NBP1-18714 |

**Supplemental Table 2. Gut microbiome characteristics. A**) Statistics of metagenomic sequencing of gut microbiome and time point. Fecal pellets obtained on days 3, 7, and 30. **B**) Gut microbiota taxonomic table characterized using the comprehensive mouse microbiota genome catalog (Kieser et al., 2022). **c**) Microbial biomarkers using logarithmic linear discriminant analysis (LDA) effect size (LEfSe) (Segata et al., 2011). Alpha threshold value for pairwise non-parametric Kruskal-Wallis test was 0.05 and threshold for the logarithmic LDA model score for discriminative features was 2.0. All-against-all comparison in multi-class analysis performed.

**Supplemental Table 3. Differential expressed genes (DEGs) between control and rapamycin group** at **A**) day 7 and **B**) day 30. Significant differential expression assessed using DEseq2 (Anders and Huber, 2010) with an FDR value ≤ 0.05 and Fold-of-change (FC)>2.

**Supplemental Table 4. Metabolome of intraluminal stool of no treatment control and rapamycin group after 7 days of treatment**. **A**) Luminal metabolites; **B**) summarized by annotated functional pathways using KEGG BRITE hierarchical classification system (Hattori et al., 2010).

**Supplemental Table 5. Gut microbiome characteristics during allogeneic stimulation. A**) Statistics of metagenomic sequencing of gut microbiome information. **B**) Gut microbiota taxonomic table characterized using the comprehensive mouse microbiota genome catalog (Kieser et al., 2022). **C**) Microbial biomarkers using logarithmic linear discriminant analysis (LDA) effect size (LEfSe) (Segata et al., 2011). Alpha threshold value for pairwise non-parametric Kruskal-Wallis test 0.05, and threshold for the logarithmic LDA model score for discriminative features 2.0. All-against-all comparison in multi-class analysis performed.

**Supplemental Tables 2-5** are available to download at https://www.dropbox.com/scl/fo/zl8ju1oiqgcembdhjekvn/AAOnxFqi6PUaeK2LWyLdoJc?rlkey=fulazk0xmaumfgbulwpye2k5t&dl=0.

